# Insights into the recruitment of the H3K4me3 reader Spp1 by the meiotic double-strand break protein Mer2

**DOI:** 10.1101/2025.09.26.678740

**Authors:** Pascaline Liloku, David Alvarez Melo, Victor G. Gisbert, Sarah Mannaert, Karen Mechleb, Sylvie Derclaye, Ankita Ray, David Alsteens, Alexander N. Volkov, Yann G.-J. Sterckx, Corentin Claeys Bouuaert

## Abstract

The formation of DNA double-strand breaks (DSBs) by Spo11 is tied to the loop-axis organization of meiotic chromosomes. Prior to DSB formation, chromatin loops marked by histone H3K4 trimethylation become tethered to the chromosome axis through interactions between Spp1 and Mer2. Mer2 is an essential partner of Spo11 thought to assemble the DSB machinery via biomolecular condensation, but these molecular assemblies remain poorly characterized. Here, using AlphaFold modeling, biochemical reconstitution, and biophysical validation, we explored the relationship between Mer2, Spp1 and their DNA-bound complexes. The tetrameric coiled-coil domain of Mer2 has four rotationally symmetrical sites that can bind a C-terminal α-helix of Spp1. However, binding of one Spp1 subunit appears to allosterically modulate the affinity for the adjacent sites, leading to the assembly of 4×2 Mer2-Spp1 complexes. Mer2 also accommodates multiple DNA duplexes, allowing the assembly of tripartite Mer2-Spp1-DNA complexes with branched DNA substrates and effective recruitment of Spp1 within nucleoprotein condensates. However, because the Spp1- and DNA-binding sites of Mer2 partially overlap, Spp1 recruitment reduces DNA binding by Mer2, which is compensated for by a patch of positively charged residues within Spp1. These findings provide insights into the structural organization of Mer2 and Spp1 and their role in the assembly of the meiotic DSB machinery.

## Introduction

During meiotic prophase I, DNA double-strand breaks (DSBs) introduced by the topoisomerase-related Spo11 protein trigger a DNA repair mechanism by recombination, which is essential for the accurate segregation of homologous chromosomes (Yadav and Claeys Bouuaert 2021; Zickler and Kleckner 2023). In *S. cerevisiae*, Spo11 acts in concert with at least nine essential partners to catalyze break formation and their activity is linked to chromosome organization (Lam and Keeney 2015). Meiotic chromosomes form linear arrays of 10-20 kb DNA loops anchored along a proteinaceous axis (Ur and Corbett 2021). Spo11 partners Rec114, Mei4 and Mer2 (RMM) are localized along the chromosome axis, but break formation happens within DNA loops (Blat et al. 2002; Pan et al. 2011; Panizza et al. 2011; Lam and Keeney 2015). Breaks are induced within 100-300 bp hotspots located in nucleosome-depleted regions flanked by nucleosomes marked by histone H3 lysine 4 trimethylation (H3K4me3) (Borde et al. 2009; Pan et al. 2011; Tischfield and Keeney 2012). The tethered loop-axis model suggests that cleavage by axis-immobilized Spo11 requires prior tethering of a DNA loop to the axis (Blat et al. 2002). Spp1 was proposed to mediate this tethering through direct interactions with H3K4me3 and axis-bound Mer2, but the underlying molecular assemblies remain poorly defined (Acquaviva et al. 2013; Sommermeyer et al. 2013).

Spp1 is a member of the Set1 complex responsible for the deposition of H3K4me3 marks (Schneider et al. 2005; Dehé et al. 2006). In the absence of Set1 or Spp1, overall DSB levels are reduced throughout the genome, although a subset of cold regions become hotter (Sollier et al. 2004; Borde et al. 2009; Acquaviva et al. 2013; Sommermeyer et al. 2013). The function of Spp1 in shaping the meiotic DSB landscape is independent of its interaction with Set1 but depends on direct binding to H3K4me3 marks and Mer2 (Acquaviva et al. 2013; Sommermeyer et al. 2013; Adam et al. 2018).

In contrast to Spp1, which is dispensable for DSB formation, the absence of Mer2 abrogates meiotic DNA breaks (Rockmill et al. 1995). In addition to binding Spp1, Mer2 also interacts with axis component Hop1 and other DSB proteins, including Rec114-Mei4, the Spo11 core complex, and the MRX complex (Arora et al. 2004; Henderson et al. 2006; Claeys Bouuaert et al. 2021; Rousová et al. 2021; Priyadarshini et al. 2025). Hence, Mer2 is thought to constitute an assembly hub for the meiotic DSB machinery.

Mer2 also has DNA-binding activity that is essential for DSB formation and undergoes DNA-dependent condensation *in vitro* (Claeys Bouuaert et al. 2021; Daccache et al. 2023). Based on these findings, Mer2, together with Rec114 and Mei4, was proposed to function as a scaffold to assemble chromatin sub-compartments (Claeys Bouuaert et al. 2021). Within those compartments, Mer2, Rec114 and Mei4 are thought to recruit Spo11 and other associated proteins, thereby controlling DSB induction (Claeys Bouuaert et al. 2021; Yadav and Claeys Bouuaert 2021; Oger and Claeys Bouuaert 2025).

Here, we investigated the interaction between Mer2 and Spp1 and their relationship with the DNA-binding and condensation activities of Mer2. Our results suggest that the recruitment of Spp1 to Mer2 tetramers is controlled allosterically to assemble complexes with a 4×2 stoichiometry. Spp1 binding partially occludes the DNA-binding surface of Mer2. However, the cost associated with Spp1 recruitment is compensated for by direct interactions of Spp1 with DNA. This work provides molecular insights into the assembly of the meiotic DSB machinery.

## Results

### Single-molecule probing of Mer2-Spp1 interaction by atomic force spectroscopy

Mer2 is a 314 amino-acid protein with a central coiled-coil domain (residues 41 to 224) flanked by N- and C-terminal intrinsically disordered regions (IDRs) (**Figure 1A**). The coiled coil assembles as a parallel tetrameric bundle, the C-terminal half of which (138-224) is necessary and sufficient for tetramerization (Daccache et al. 2023) and contains the Spp1 interaction domain (Acquaviva et al. 2013) (**Figure 1B**). Mutation of a conserved V195 residue within Mer2, predicted to be surface exposed, abolishes Mer2-Spp1 co-immunoprecipitation and recapitulates the meiotic phenotype of a *spp1Δ* strain (Adam et al. 2018).

**Figure 1:**
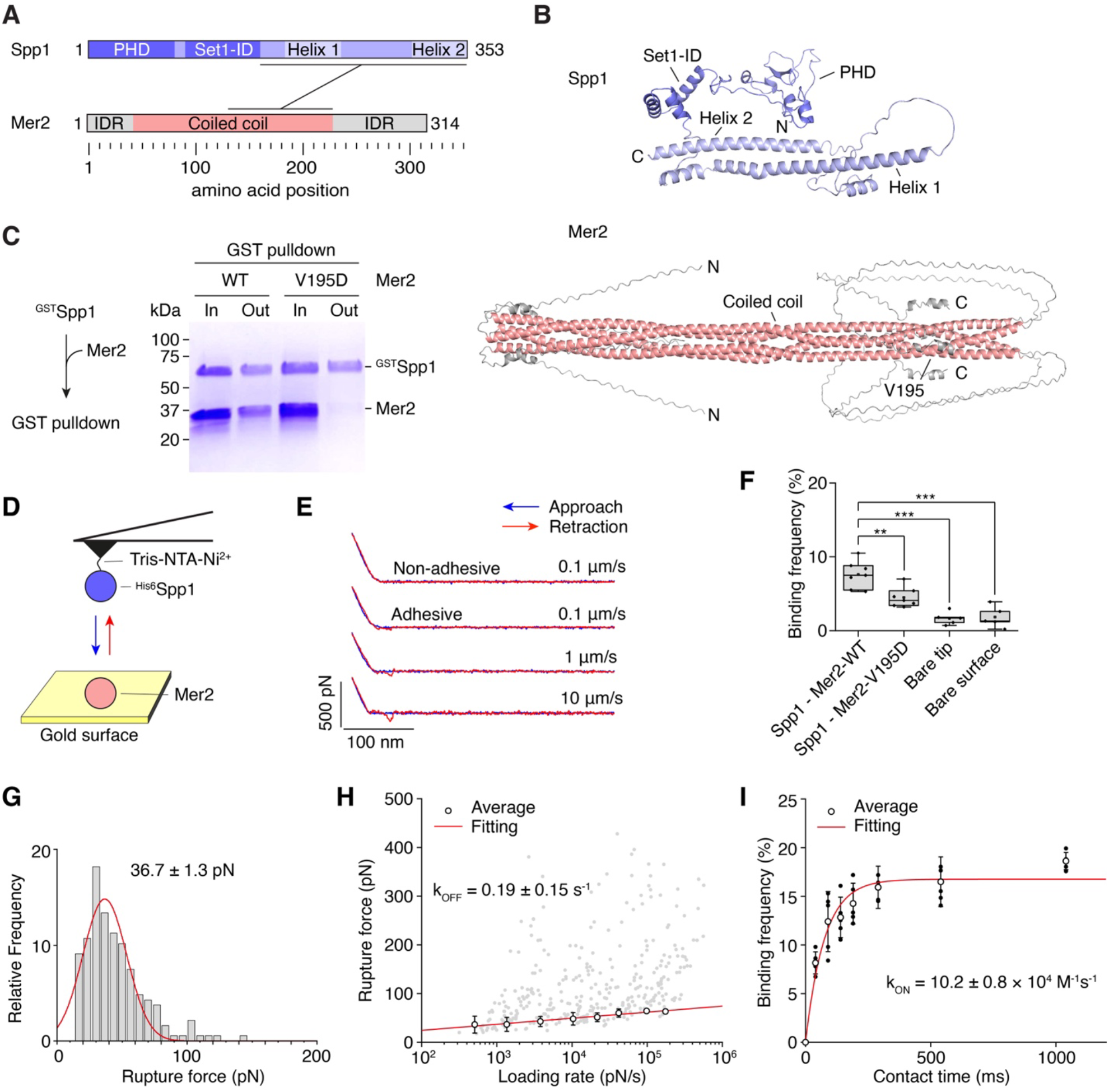
Single-molecule analysis of the interaction between Mer2 and Spp1. **(A)** Domain structure Spp1 and Mer2. PHD, plant homeodomain; Set1-ID, interaction domain; IDR, intrinsically disordered region. The Mer2-Spp1 interaction domain is indicated. **(B)** AlphaFold predictions of Spp1 (AF-Q03012-F1) and tetrameric Mer2. The position of the Spp1-interacting residue V195 is shown. **(C)** Pulldown of wild-type and mutant (V195D) Mer2 with GST-tagged Spp1. In, input; Out, output. **(D)** Setup of the AFM assay to quantify the kinetics and thermodynamics of the interaction between Spp1 and Mer2. **(E)** Examples of force-distance curves obtained upon approach and retraction of the tip from the surface at different loading rates. **(F)** Binding frequency between tip-immobilized Spp1 and surface-bound wild-type or mutant (V195D) Mer2, compared to controls without Spp1 or Mer2. Mean ± SD of n = 5-7 independent experiments. **(G)** Histogram of the forces measured upon tip retraction (range of loading rate 350-900 pN/s). The data was fitted to a normal distribution (R^2^ = 0.88). **(H)** Dynamic force spectroscopy analysis showing the rupture force as a function of loading rate for individual measurements (grey dots indicate individual unbinding events, n = 6 experiments). Open circles represent the mean rupture force obtained by fitting histograms at discrete loading rates and error bar represent the S.D. (see **Supplementary Figure 2**). The red solid line represents the Bell-Evans fit extrapolated to zero-force. **(I)** Binding frequency as a function of contact time. Data points show the mean binding frequency ± S.D. from 5 adhesion maps, each consisting of 256 individual force-distance curves. Red solid line shows a non-linear curve fitted to the data.

Spp1 is a 353 amino-acid protein with an N-terminal plant homeodomain (PHD) finger that binds H3K4me3 modifications (He et al. 2019) (**Supplementary Figure 1A**), a Set1 interaction domain (Set1-ID) (**Supplementary Figure 1B**), and a C-terminal region predicted to contain two long α-helices, here referred to as helix 1 and 2, required for Mer2 binding (Acquaviva et al. 2013) (**Figure 1A, B**).

To verify that the *in vitro* interaction between Spp1 and Mer2 depends on Mer2-V195, as shown *in vivo* (Adam et al. 2018), we incubated GST-tagged Spp1 with wild-type Mer2 or a V195D mutant and pulled down proteins by glutathione affinity. SDS-PAGE analysis showed that ^GST^Spp1 efficiently interacts with wild-type Mer2, but not the V195D mutant (**Figure 1C**).

We set up a single-molecule interaction assay based on atomic force microscopy (AFM) (Müller et al. 2021; Fernandez et al. 2023) to further characterize the interaction between Mer2 and Spp1. Spp1 was grafted to an Ni^2+^-NTA coated tip through an N-terminal His_6_ affinity tag, and Mer2 was covalently immobilized to the surface by NHS-EDC chemistry (**Figure 1D**) (Alsteens et al. 2015). The interaction between Spp1 and Mer2 was measured by bringing the tip into contact with the surface and retracting it. Each approach-retraction cycle yields a force-distance (FD) curve, a subset of which reveals a binding event, detected by the deflection of the tip upon retraction (**Figure 1E**). In the presence of Spp1 and Mer2, 7.6 ± 1.8% (Mean ± S.D., N=7), of the curves showed interaction (**Figure 1F**). When Mer2 was replaced by the V195D mutant, the binding frequency dropped to 4.5 ± 1.3%. When either Spp1 was omitted from the tip, or Mer2 was omitted from the surface, the binding frequency was further decreased to about 2%. Hence, the assay captures specific single-molecule interactions between Spp1 and Mer2.

We extracted the force required to disrupt Spp1 and Mer2 interaction (rupture force), which revealed a peak at about 36.7 ± 1.3 piconewton (pN) (Mean ± S.E.M.) at a loading rate of 500 pN/s (**Figure 1G**). To determine kinetic parameters of the interactions, we measured the rupture force at different loading rates (*i*.*e*. the force acting on the interaction over time). Plotting the rupture force against the loading rate allows us to extract the free energy landscape under non-equilibrium conditions through application of a Bell-Evans model and deduce the dissociation rate (k_OFF_) (Alsteens et al. 2015). This revealed a k_OFF_ = 0.19 ± 0.15 s^−1^ (**Figure 1H, Supplementary Figure 2, Supplementary Table 1**). In addition, varying the contact time (*t*) allows us to estimate the interaction time, and thereby deduce the kinetic on-rate (k_ON_), estimated at 10.2 ± 0.8 × 10^4^ M^−1^·s^−1^ (**Figure 1I**). Hence, a single Spp1 monomer binds to Mer2 tetramers with a dissociation constant (K_D_) of 1.9 ± 1.6 µM. This corresponds to a binding affinity about two orders of magnitude lower than the ensemble average affinity previously obtained by microscale thermophoresis (MST) (Rousová et al. 2021), suggesting that the single-molecule interaction we characterized by AFM is rather different from that captured in bulk biochemical assays (see below).

### Structural model of a minimal Mer2-Spp1 complex

Previous biochemical analyses indicated that Mer2 and Spp1 form a complex with a 4×2 (Mer2:Spp1) stoichiometry (Rousová et al. 2021). To gain further insights, we used AlphaFold3 (Abramson et al. 2024) to predict the structure of a complex composed of Mer2 residues 138-224 and Spp1 residues 161-353. AlphaFold produced high-confidence models showing two Spp1 subunits bound to the surface of the Mer2 tetrameric coiled coil (**Figure 2A, Supplementary Figure 3A**).

**Figure 2:**
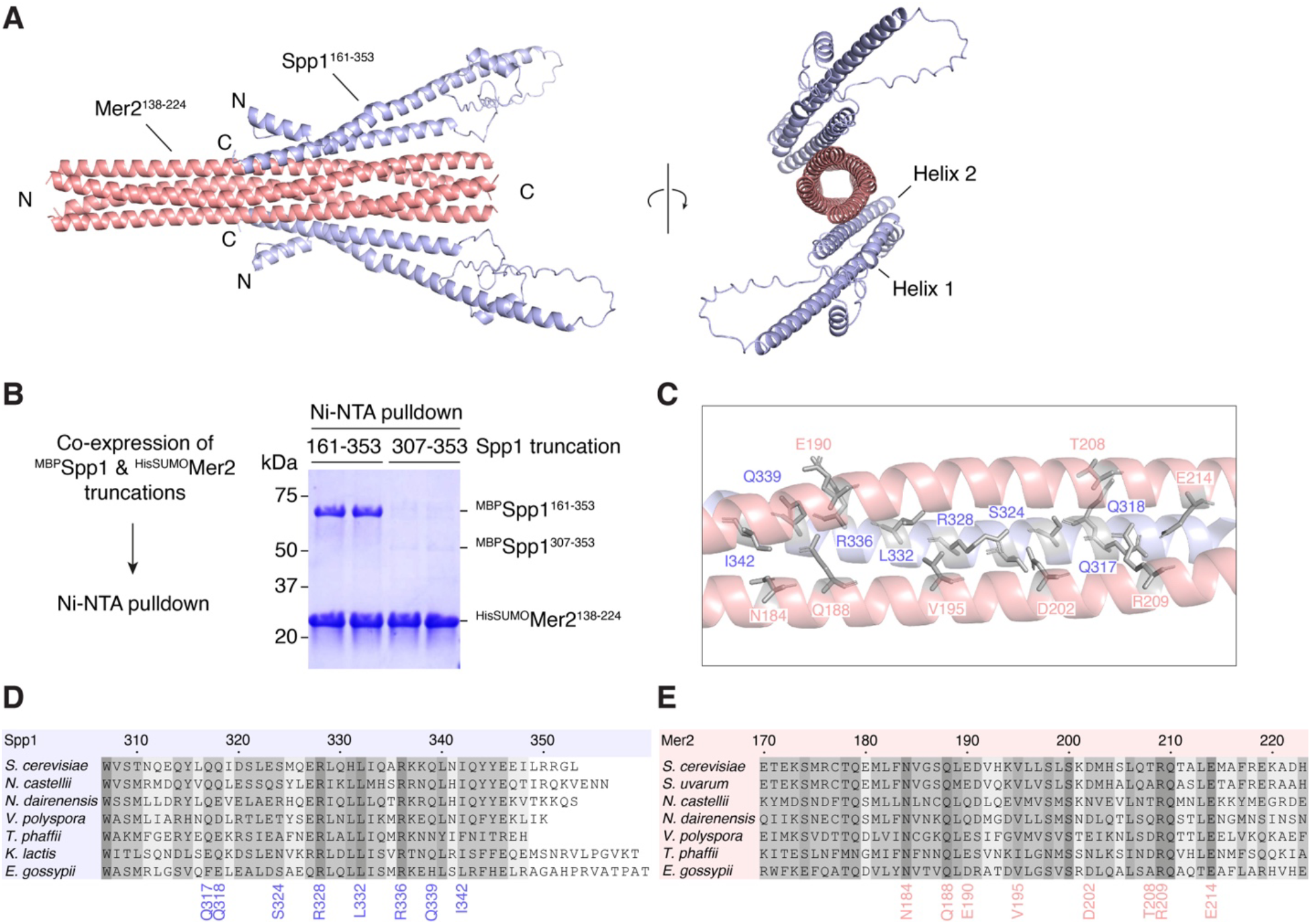
Model of a minimal Mer2-Spp1 complex. **(A)** AlphaFold prediction of a 4×2 complex between Mer2^138-224^ and Spp1^161-353^. **(B)** Pulldown of MBP-tagged Spp1 truncations with HisSUMO-tagged Mer2^138-224^. **(C)** Zoom of the interface between Mer2 and Spp1 with residues of interest highlighted. Mer2, pink, Spp1, blue. **(D, E)** Multiple sequence alignment Spp1 (D) and Mer2 (E) with residues mutated in **Figure 3** highlighted.

Within the model, the C-terminal α-helix of Spp1 (helix 2, residues W307 to R351) is bound anti-parallel to the Mer2 coiled coil. A long α-helix (helix 1, residues D169 to S233, interrupted by a kink at R181 and D182) folds back onto helix 2. In a pulldown assay, Spp1^161-353^ was sufficient for the interaction with Mer2^138-224^, but Spp1^307-353^ was not (**Figure 2B**), suggesting that helix 1 contributes to the stability of the Mer2-Spp1 interface. Consistently, in the absence of helix 1, AlphaFold could not accurately model the structure of the Mer2-Spp1 complex (**Supplementary Figure 3B**).

In the AlphaFold model, Spp1 helix 2 interacts with two adjacent Mer2 helices (**Figure 2C**). Spp1-Q317 contacts Mer2-R209 from one subunit and E214 from the other subunit; Spp1-Q318 contacts Mer2-T208; Spp1-S324 and R328 point inside the trimeric interface and contact Mer2-D202; Spp1-L332 contacts Mer2-V195; Spp1-R336 makes electrostatic interactions with Mer2-E190; Spp1-Q339 is in proximity to Mer2-Q188 and Spp1-I342 points towards Mer2-N184.

Sequence alignments of Spp1 and Mer2 homologs from seven species of the Saccharomycetaceae family indicate that most of the residues within the predicted Mer2-Spp1 interface are conserved (**Figure 2D, E**). In particular, Spp1 residues R328, L332, R336, and Mer2 residues N184, Q188, R209 and E214 are invariant in the aligned sequences. Most other residues are also conserved or replaced by similar amino acids, including Mer2-V195 that is substituted to isoleucine in *T. phaffii*. Consistent with the structural conservation of the interface, AlphaFold yielded similar models across pairs of Mer2-Spp1 homologs from Saccharomycetaceae (**Supplementary Figure 4**).

To test the AlphaFold model experimentally, we mutated the predicted interface residues to alanine and quantified the effect of the mutation on the interaction between Spp1 and Mer2 by pulldown. Wild type and mutant ^MBP^Spp1^161-353^ and ^HisSUMO^Mer2^138-224^ were co-expressed in *E. coli* and pulled down by Ni-NTA affinity. The Mer2-V195D mutant was used as a control. As expected, Mer2-V195D and V195A strongly compromised the interaction with Spp1 (**Figure 3A**). In addition, alanine substitution of Mer2 residues E190 and R209 significantly decreased the interaction with Spp1. On the Spp1 side, R328A and L332A mutations strongly decreased the interaction with Mer2, and S324A, R336A and I342A had milder defects (**Figure 3B**). Interestingly, the mutations that led to the strongest phenotypes were located towards the center of the predicted interface (**Figure 2C**). Hence, the mutagenesis analysis supports the AlphaFold model of the Mer2-Spp1 interaction domain.

**Figure 3:**
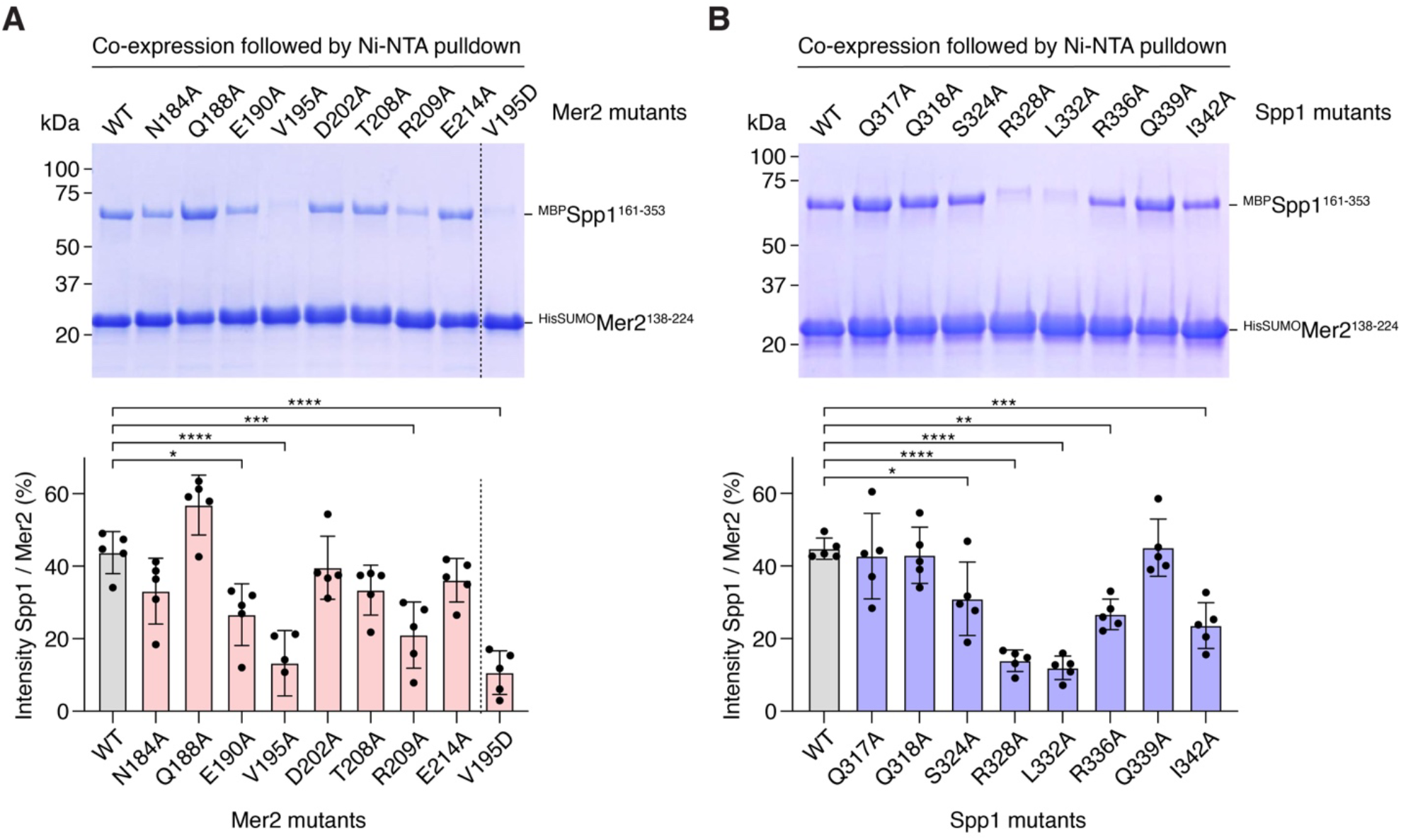
Mutational analysis of the Mer2-Spp1 interaction surface. **(A, B)** Ni-NTA pulldown analysis of MBP-tagged Spp1^161-353^ with HisSUMO-tagged Mer2^138-224^. Panel A show Mer2 mutants, panel B show Spp1 mutants. Quantifications show the mean ± SD from five independent experiments. * *p* < 0.05, ** *p* < 0.01, *** *p* < 0.001, **** < 0.0001 (one-way ANOVA).

### The stoichiometry of Mer2-Spp1 complexes

The AlphaFold model does not readily account for the reported 4×2 stoichiometry of the Mer2-Spp1 complex (Rousová et al. 2021). Indeed, given the four-fold radial symmetry of Mer2, one might expect Mer2 tetramers to accommodate up to four Spp1 subunits. Consistent with that idea, AlphaFold produces high-confidence 4×4 models without steric clashes between Spp1 subunits (**Supplementary Figure 5A**). The binding sites are identical between 4×2 and 4×4 models, and the respective position of two Spp1 subunits to adjacent or opposite sides of the Mer2 coiled coil appears to be arbitrary (**Supplementary Figure 5B**).

To clarify this, we sought to revisit the stoichiometry of Mer2-Spp1 complexes. We co-expressed MBP-tagged Spp1 and HisSUMO-tagged Mer2, purified a complex by Ni-NTA affinity chromatography and analyzed purified complexes by size-exclusion chromatography followed by multi-angle light scattering (SEC-MALS). This yielded a molecular mass of 365 ± 6 kDa, consistent with a 4×2 stoichiometry (expected MW = 370 kDa). We also produced a collection of tagged and untagged full-length and truncated complexes and analyzed the samples by SEC-MALS (**Supplementary Figure 6**) and small angle X-ray light scattering (SAXS) (**Supplementary Table 2**). All the configurations yielded a 4×2 stoichiometry, regardless of the analysis method (**Figure 4A**).

**Figure 4:**
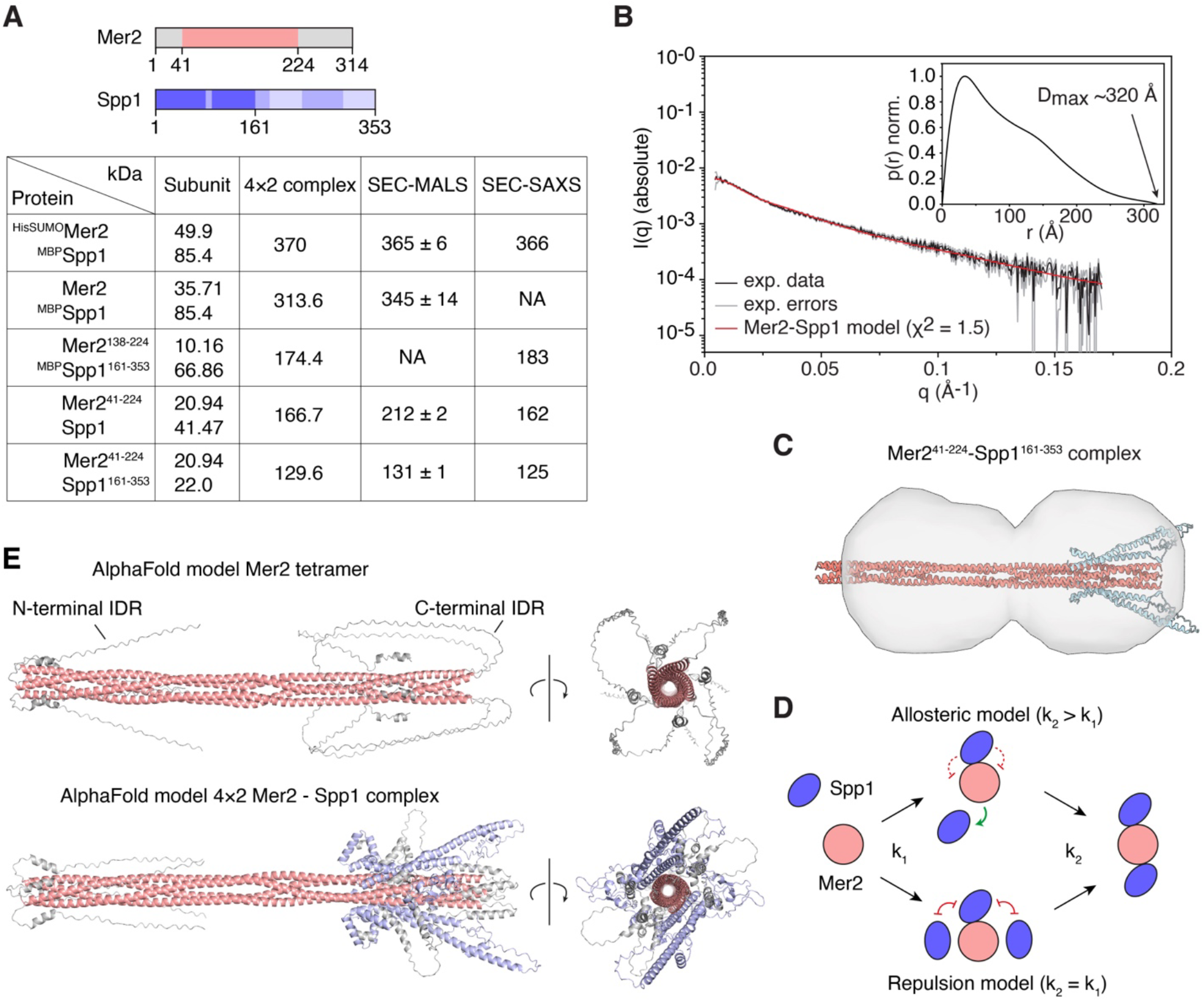
SEC-MALS and SAXS analyses of Mer2-Spp1 complexes. **(A)** Molecular weight measurements of Mer2-Spp1 complexes by SEC-MALS and SEC-SAXS (see **Supplementary Figure 7 and Supplementary Table 2**). **(B)** SEC-SAXS analysis of Mer2^41-224^-Spp1^161-353^ complexes. The main graph shows the experimental data (black), error margins (gray), and the fit of the AlphaFold model to the data (red). The inset shows a normalized probability distance distribution function obtained from the experimental SAXS data, with the approximate D_max_ value indicated. **(C)** Overlay of Mer2-Spp1 model with the *ab initio* reconstructed shape. **(D)** Allosteric and repulsion models can explain the preferential binding of two Spp1 subunits on opposite binding sites of the Mer2 coiled coil. The cartoon illustrates a cross-section perpendicular to the coiled coil axis. **(E)** AlphaFold models of full-length Mer2 tetramer and 4×2 Mer2-Spp1 complexes. The coiled-coil domain of Mer2 is colored in salmon, the IDRs are grey, Spp1 is blue.

We considered the possibility that the purification protocol enriches for Mer2 compared to Spp1 subunits. However, when Spp1 was added in excess to Mer2, the complexes still yielded a 4×2 stoichiometry (**Supplementary Figure 6C, D**). Hence, Mer2 tetramers only accommodate two Spp1 subunits and the underlying reason is not immediately evident from the AlphaFold models.

To further test the AlphaFold models, we used SAXS to determine the overall shape of the Mer2-Spp1 complexes. SAXS measurements of a 4×2 Mer2^41-224^-Spp1^161-353^ complex were in excellent agreement with the curve expected based on structural models showing two Spp1 subunits bound to opposite sides of the Mer2 tetrameric coiled coil (×^2^ = 1.5, **Figure 4B**) and *ab initio* particle reconstruction from SAXS data were also consistent with the model (**Figure 4C**). Similarly, a complex containing the Mer2 coiled coil and full-length Spp1, or a shorter truncation with MBP-tagged Spp1 were all consistent with AlphaFold models (**Supplementary Figure 7**).

We consider two main scenarios that can account for the observed 4×2 stoichiometry: either Spp1 subunits cannot occupy adjacent binding sites on Mer2, perhaps due to steric clashes or charge repulsions, or binding of Spp1 induces allosteric regulation that causes positive cooperativity between opposite sites, and perhaps negative cooperativity between adjacent sites (**Figure 4D**). The latter model would explain the low binding affinity obtained with our AFM assay, compared to the ones previously obtained by MST (Rousová et al. 2021). Indeed, our assay captures single-molecule interactions that, under this scenario, would correspond to a low-affinity primary contact between Spp1 and Mer2.

### The full-length Mer2-Spp1 complex

To gain further insights into the Mer2-Spp1 complex, we used AlphaFold to model the structure of a full-length Mer2 tetramer and a 4×2 Mer2-Spp1 complex (**Figure 4E**). AlphaFold consistently placed two Spp1 subunits on opposite sides of the Mer2 α-helical bundle, through the same interaction surface as for the truncated versions presented above. The position of the Spp1 PHD finger and Set1-ID varied between models (**Supplementary Figure 8**) and the C-terminal Mer2 IDR was placed loosely around interaction domain.

To address whether the positioning of the Mer2 C-terminal IDR in these models was relevant, we asked whether the IDR contributes to the interaction between Mer2 and Spp1. By pulldown analysis, we found that the absence of the Mer2 C-terminal IDR led to a small but reproducible reduction in Spp1 binding (**Supplementary Figure 9A**). AFM analysis also revealed a slight reduction in the binding frequency between Spp1 and Mer2^1-224^, compared to the full-length control (**Supplementary Figure 9B**), suggesting that the C-terminal Mer2 IDR may weakly contribute to the interaction.

### The tripartite Mer2-Spp1-DNA complex

We previously showed that Mer2 binding to DNA involves both the Mer2 coiled-coil domain and the C-terminal IDR (Claeys Bouuaert et al. 2021; Daccache et al. 2023). Indeed, *in vitro*, the coiled-coil domain is necessary and sufficient for DNA binding. However, mutating IDR residues K265, R266, R267 and R268 (KRRR) to alanine severely compromises DNA-binding and condensation of Mer2 *in vitro* and abolishes meiotic DSB formation *in vivo* (Claeys Bouuaert et al. 2021).

In the absence of N- and C-terminal IDRs, Mer2 does not assemble condensates *in vitro*, but retains a DNA-bridging activity and binds preferentially to branched DNA substrates compared to duplex DNA, indicating that Mer2 can bind simultaneously to multiple DNA duplexes (Daccache et al. 2023). Based on AFM imaging and comparisons with Mer2 orthologs (*S. pombe* Rec15 and *M. musculus* IHO1), we previously proposed that the coiled-coil domain binds co-aligned DNA duplexes through electrostatic interactions spread along the length of the coiled coil (Daccache et al. 2023). Interestingly, AlphaFold proposed a structural model of DNA-bound Mer2 tetramers entirely consistent with those expectations, with DNA duplexes aligned parallel to the coiled coil axis and the KRRR motif contacting the DNA double helix (**Figure 5A**).

**Figure 5:**
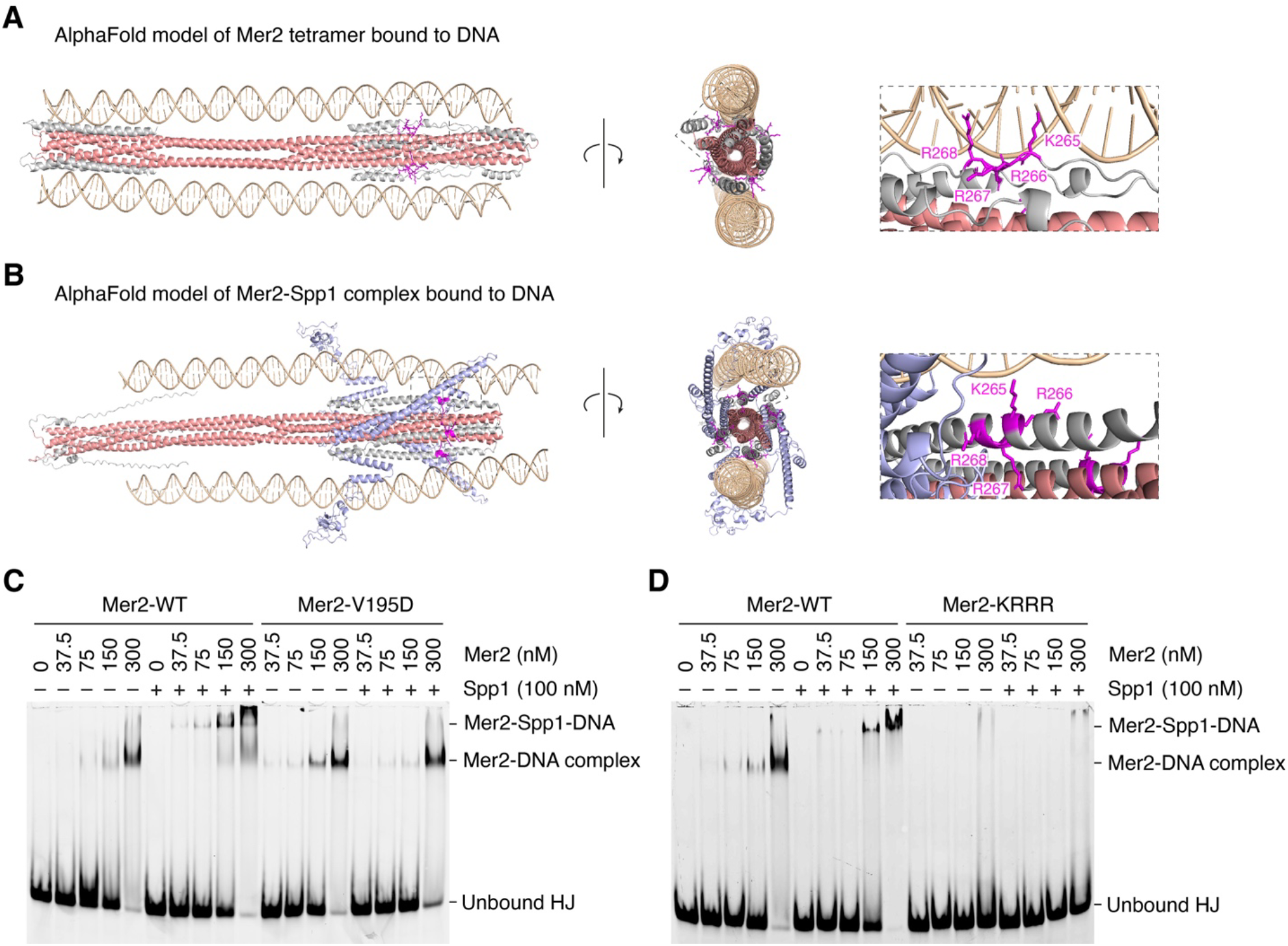
The tripartite Mer2-Spp1-DNA complex. **(A, B)** AlphaFold3 models of full-length Mer2 and Mer2-Spp1 complexes bound to two molecules of 80 bp double-stranded DNA. The position of the Mer2 DNA-binding residues (KRRR) is shown (magenta). **(C, D)** Formation of a tripartite complex between wild type or mutant Mer2, Spp1 and a branched DNA substrate (HJ, Holliday Junction with 20 bp arms).

Since the coiled-coil domain of Mer2, in particular the C-terminal half (residues 138-224), is both involved in binding Spp1 and DNA, we wondered how the presence of Spp1 and DNA influence each other and whether both binding partners could be accommodated simultaneously.

To address this, we modeled a 4×2 Mer2-Spp1 complex in the presence of two molecules of duplex DNA. Similar to the model without Spp1, the DNA duplexes were modeled parallel to the Mer2 coiled coil with the Mer2-KRRR motif bound to DNA (**Figure 5B**).

Next, we sought to reconstitute a tripartite Mer2-Spp1-DNA complex *in vitro*. To do this, we performed gel shift analyses using a four-way branched DNA substrate (Holliday Junction, HJ) assembled with partially overlapping 40 nt oligonucleotides. In the presence of Spp1, we observed a complex of lower electrophoretic mobility compared to the Mer2-DNA complex formed in the absence of Spp1 (**Figure 5C**). When reactions contained the Mer2-V195D mutant, the assembly of the tripartite Mer2-Spp1-DNA complex was abolished, but the Mer2-DNA complex was not affected. In addition, when the reactions contained the Mer2-KRRR mutant, both the Mer2-DNA and Mer2-Spp1-DNA complexes were absent (**Figure 5D**).

Quantification of Mer2 titrations indicates that the presence of Spp1 mildly stimulates DNA binding (**Supplementary Figure 10A**). Hence, despite occupying similar binding sites, we observe no competition between Spp1 and DNA for access to Mer2. Consistently, in an order-of-addition experiment, the sequence in which Mer2 was incubated in the presence of Spp1 and DNA did not significantly impact the assembly of the complexes, further indicating that complex formation reached equilibrium before electrophoresis (**Supplementary Figure 10B**). This equilibrium was reached very rapidly, since Mer2-Spp1-DNA complexes were formed efficiently even if Spp1 was added just before loading the gel (**Supplementary Figure 10C**).

### Spp1 is recruited to Mer2 condensates as a client

During meiosis, Spp1 has separable functions as part of the Set1 complex and as a Mer2 binding partner (Adam et al. 2018). The diffusion rate of Spp1 is lower when it is associated with Mer2 compared to the Set1 complex (Karányi et al. 2018), which can be explained by the recruitment of Spp1 within Mer2 condensates (Claeys Bouuaert et al. 2021).

To investigate the relationship between Mer2 condensation and Spp1 recruitment, we mixed eGFP-tagged Mer2 and mScarlet-tagged Spp1 in the presence of plasmid DNA. This led to the formation of bright fluorescent foci with essentially perfect overlap between the two colors (**Figure 6A**). When condensates were assembled with the V195D mutant, Mer2 foci formed efficiently but the Spp1 fluorescent signal was absent (**Figure 6B**). While Spp1 is not required for the formation of Mer2 foci (**Figure 6C**), Mer2 is necessary for efficient formation of Spp1 foci (**Figure 6D**). Nevertheless, Spp1 slightly stimulates Mer2 condensation. Hence, Spp1 is recruited to Mer2 foci essentially as a client, but it seems to contribute to DNA binding and condensation.

**Figure 6:**
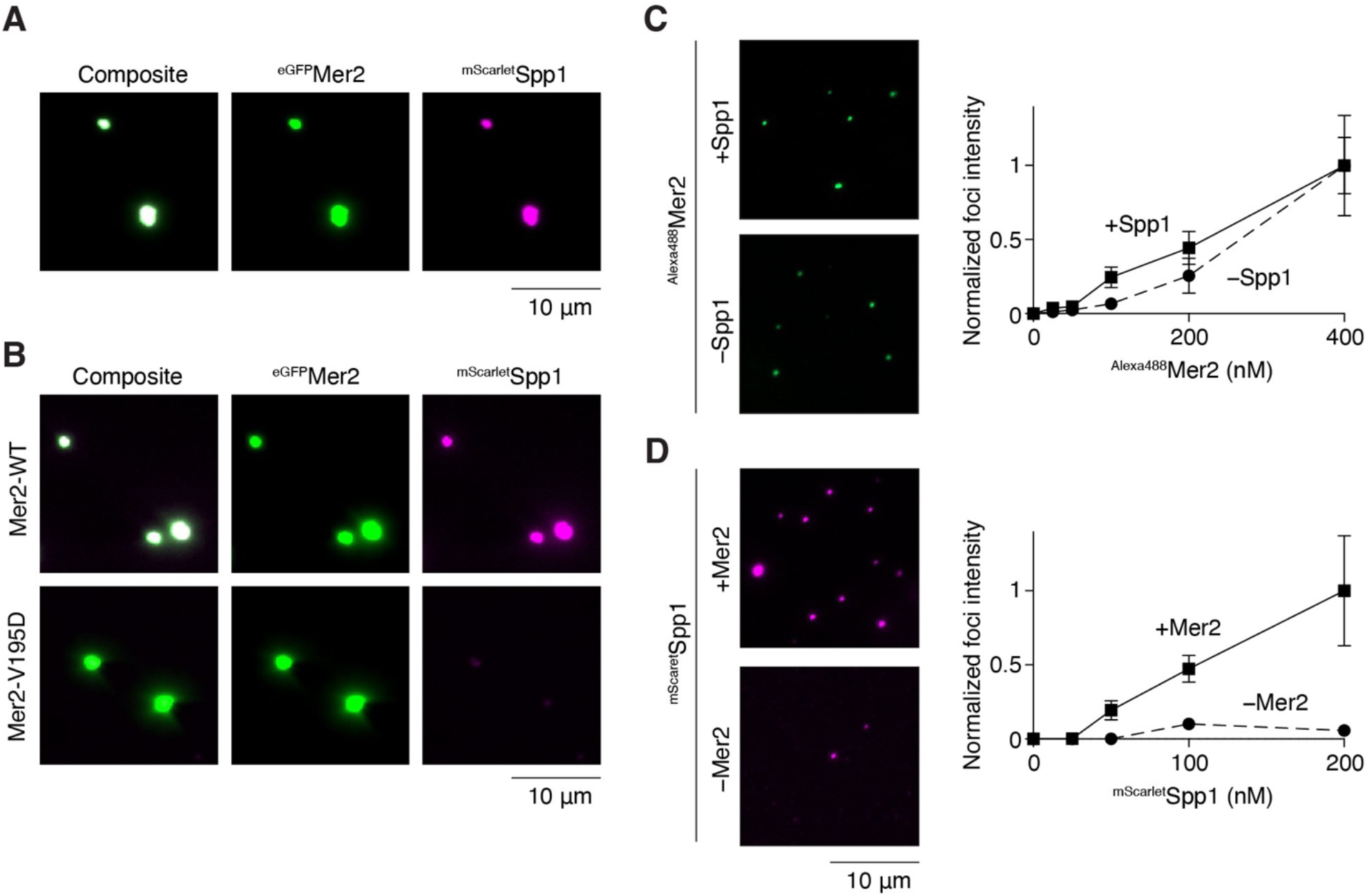
Mer2-Spp1 nucleoprotein condensates. **(A)** Visualization of nucleoprotein condensates by epifluorescence microscopy using tagged Mer2 and Spp1. **(B)** Effect of the Mer2-V195D mutation on the colocalization of Mer2 and Spp1 condensates. **(C)** Impact of Spp1 on the formation of Mer2 foci. Quantifications show the average focus intensity (normalized to 400 nM Mer2-Spp1) **(D)** Impact of Mer2 on the formation of Spp1 foci. Quantifications show the average focus intensity (normalized to 200 nM Spp1 + Mer2). Error bars show mean ± SD from ten fields of view.

### Spp1 directly contributes to DNA binding

We sought to address whether the weak stimulatory activity of Spp1 on Mer2 DNA binding and condensation are caused by direct interaction of Spp1 with DNA. We first asked whether Spp1 has DNA-binding activity, but a gel-shift analysis provided no evidence for DNA binding by full-length Spp1 (**Figure 7A**). However, the C-terminal domain of Spp1 alone (residues 161-353) bound DNA quite efficiently (**Figure 7A**). This suggests that the Spp1 N-terminal domain occludes a DNA-binding site within Spp1.

**Figure 7:**
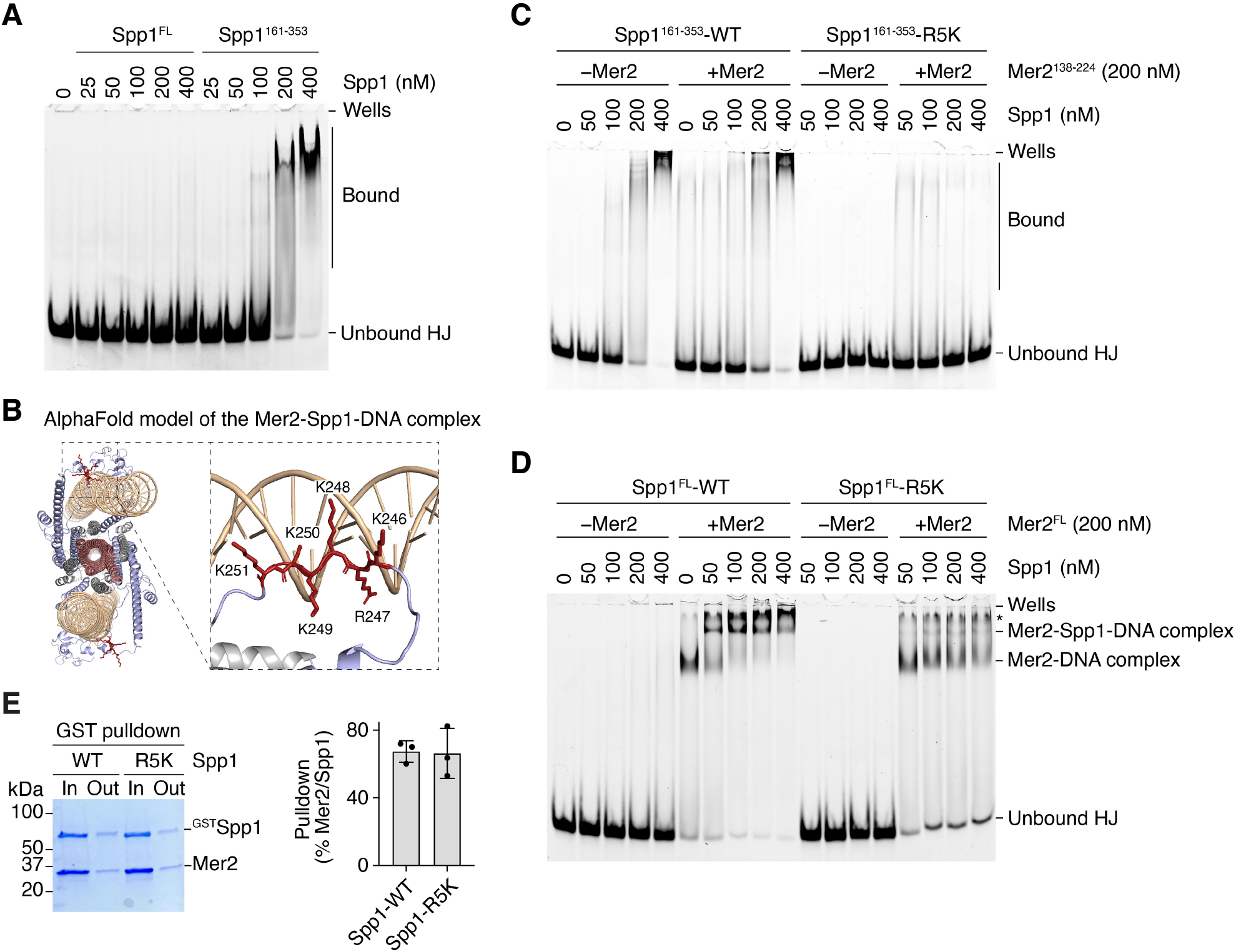
DNA-binding activity of Spp1. **(A)** Gel shift assay of full-length or truncated Spp1 binding to a branched DNA substrate (HJ, Holliday Junction with 20 bp arms). (**B)** AlphaFold3 model of Mer2-Spp1-DNA complexes. The position of the Spp1-R5K motif is highlighted. **(C)** Gel shift assay of wild-type (WT) and mutant (R5K) Spp1^161-353^ truncations on a HJ substrate in the presence or absence of Mer2^138-224^ truncation. The Spp1-R5K mutant has residues K246, R247, K248, K249, K250, K251 mutated to alanine. **(D)** Gel shift assay of wild-type and mutant full-length Spp1 DNA binding in the presence or absence of full-length Mer2. The band labeled * corresponds to a higher-order Mer2-(Spp1)-DNA assembly, which we interpret as reflecting the condensation property of Mer2. The detection of this band varied somewhat between experiments. **(E)** Pulldown of Mer2 with wild-type and mutant (R5K) GST-tagged Spp1. In, input; Out, output. Quantifications show the mean ± SD from three replicates.

In the AlphaFold model of the Mer2-Spp1-DNA complex, a patch of positively charged residues located between helix 1 and helix 2 (K246, R247, K248, K249, K250, K251, here referred to as R5K) is shown to interact with DNA (**Figure 7B**). To test whether these residues are responsible for the DNA-binding activity of Spp1, we mutated this stretch of residues to alanine and established the impact of the mutation on DNA binding by a gel shift assay. In the context of truncations that only included the minimal Mer2-Spp1 interaction domains, Mer2^138-224^ alone bound very weakly to the HJ substrate and Spp1^161-353^ strongly promoted DNA binding. As expected, the Spp1-R5K mutation completely abolished this stimulatory effect (**Figure 7C**).

In the context of full-length proteins, wild-type Spp1 did not significantly bind DNA by itself but enabled the assembly of a tripartite complex in the presence of Mer2. Mutation of the R5K motif strongly decreased the assembly of the tripartite complex and Mer2-DNA complexes were formed instead (**Figure 7D**). The transition from Mer2-Spp1-DNA to Mer2-DNA complex could be explained if the R5K mutation affects Mer2-Spp1 interaction. However, this was not the case (**Figure 7E**).

These results show that, even though full-length Spp1 does not bind DNA by itself, it does so in the context of a complex with Mer2. However, mutating a DNA-binding site within a complex is expected to reduce the overall affinity of the complex for DNA. This is not what we observed, and the Spp1-R5K mutation instead drove the formation of Mer2-DNA complexes at the expense of Mer2-Spp1-DNA assemblies, suggesting an alternative explanation. We propose that the presence of Spp1 inherently reduces the DNA-binding activity of Mer2, which is counter-balanced by binding of the Spp1-R5K motif to DNA. When the R5K motif is mutated, the cost associated with Spp1 binding to Mer2 drives the equilibrium in favor of free Mer2 complexes (**Figure 8A**).

**Figure 8:**
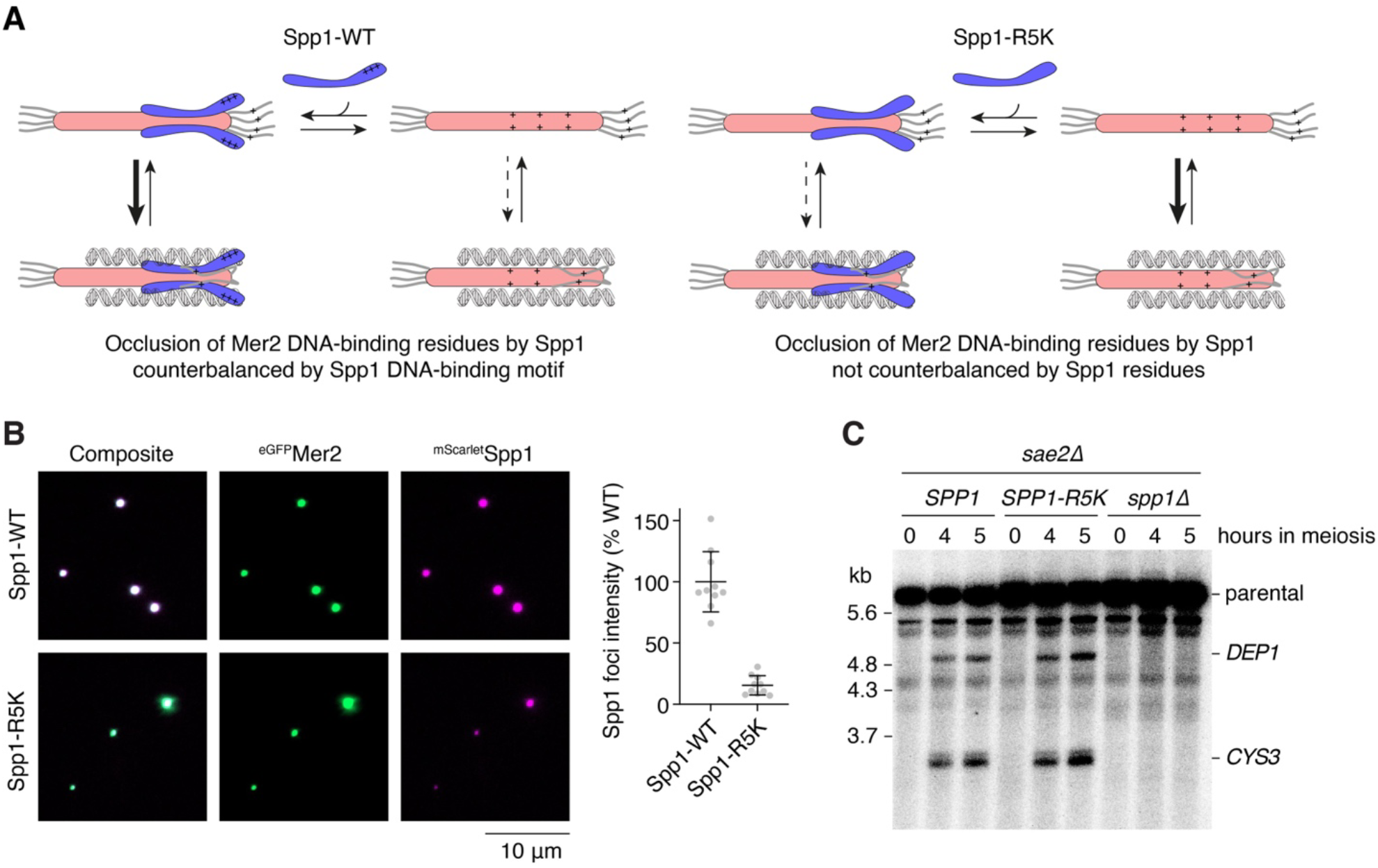
Interplay between Mer2, Spp1 and DNA binding. **(A)** Model for the role of Spp1 DNA binding function. (Left) We propose that Spp1 binding to Mer2 occludes DNA-binding residues (+) within Mer2. This is compensated by the Spp1-R5K motif that drives the equilibrium in favor of Mer2-Spp1-DNA complexes. (Right) When the R5K motif is mutated, the decreased DNA-binding activity of Mer2 caused by Spp1 binding drives the equilibrium to Mer2-DNA complexes. **(B)** Effect of Spp1-R5K mutation on the recruitment of Spp1 within Mer2 condensates. Quantifications show the total focus intensity per field of view, normalized to Spp1-WT. Error bars are mean ± SD (n = 10 fields of view). **(C)** Southern blot analysis of meiotic DSB formation at the *CYS3* and *DEP1* hotspots in wild-type and Spp1 mutant strains.

A prediction of this model is that the Spp1-R5K mutant would be excluded from Mer2 condensates, even though the protein-protein interaction between Mer2 and Spp1 is unaffected. Consistently, we found that mutation of the R5K motif severely reduced the recruitment of Spp1 to Mer2 condensates *in vitro* (**Figure 8B**).

*In vivo*, the absence of Spp1 leads to a severe reduction of DSB levels at some hotspot (e.g. *CYS3* and *DEP1*) and an increase in DSBs at others (e.g. *PES4*) (Sommermeyer et al. 2013). We sought to evaluate the relevance of the Spp1 DNA-binding motif in shaping the meiotic DSB landscape. Since the R5K mutation leads to exclusion of Spp1 from Mer2 condensates *in vitro*, we hypothesized that the mutation may recapitulate the phenotype of a *spp1Δ* strain, even though the H3K4me3 and Mer2-binding functions of Spp1 are unaffected. However, this was not the case. Southern blot analysis of DSB formation in repair-defective *sae2Δ* or *dmc1Δ* backgrounds indicated that the *SPP1-R5K* mutation does not reduce DSB formation at *CYS3* and *DEP1* hotspots, and does not stimulate DSB formation at the *PES4* hotspot, in contrast to the *spp1Δ* mutation (**Figure 8C, Supplementary Figure 11**). Hence, although the R5K motif promotes DNA binding by Spp1 *in vitro*, it is not required for Spp1 meiotic function *in vivo*.

## Discussion

Here, we used biochemical reconstitution, single-molecule analyses, and AlphaFold modeling to explore the interaction between Mer2 and Spp1 and their relationship with DNA. We determined thermodynamic and kinetic parameters of the Mer2-Spp1 interaction and presented structural models of *S. cerevisiae* Mer2-Spp1 complexes, supported by SAXS and mutagenesis. In addition, we show that Spp1 is recruited to Mer2 condensates and identified a motif within Spp1 that fosters the interaction of Mer2-Spp1 complexes with DNA. Our results have implications for the role of Mer2 and Spp1 in the assembly of the meiotic DSB machinery.

### Assembly of the Mer2-Spp1 complex

Consistent with previous research (Rousová et al. 2021), we have shown that Mer2 and Spp1 assemble a complex with a 4×2 stoichiometry, where the C-terminal α-helix of Spp1 (helix 2, residues 307-351) binds anti-parallel to and interacts with two adjacent helices at the C-terminus of the Mer2 coiled coil (residues 180-224). Spp1 helix 1 (residues 169-233) folds on top of helix 2 and stabilizes this interface. This model is supported by pulldown assays and mutagenesis, and is consistent with previous truncation (Acquaviva et al. 2013) and mutagenesis (Adam et al. 2018) experiments in yeast.

Nevertheless, the structural model proposed by AlphaFold does not readily account for the 4×2 stoichiometry of the complex. Instead, the model suggests that the tetrameric coiled-coil domain of Mer2 has four binding sites for Spp1 that could be occupied simultaneously. It is possible that some aspects of the model are inaccurate and four Spp1 subunits would sterically clash or else repulse each other. However, we favor an alternative explanation, based on allosteric modulation of the Mer2-Spp1 interaction domain. We suggest that binding of one Spp1 subunit to Mer2 induces tension on the coiled coil that increases the affinity of the opposite Spp1-binding site, and perhaps reduces the affinity of the two adjacent sites (**Figure 5E**). In other words, binding of Spp1 is positively cooperative at opposite sides of the coiled coil, but negatively cooperative at adjacent sites, leading to the symmetrical arrangement of two Spp1 subunits on opposite sides of the Mer2 coiled coil. Evidence supporting this model comes from our AFM-based force spectroscopy assay that revealed a binding affinity of single Spp1 monomers to Mer2 tetramers about 100x lower than those obtained from ensemble average experiments (Rousová et al. 2021). In addition, this allosteric model would also explain why varying the ratio of Mer2 and Spp1 in a gel shift assay only reveals a single Mer2-Spp1-DNA complex rather than multiple complexes of distinct stoichiometries. Nevertheless, elucidating this mechanism will require high-resolution structural determination of Mer2-Spp1 complexes.

### Assembly of the Mer2-Spp1-DNA complexes

AlphaFold modeling suggested that Mer2 tetramers can bind co-aligned DNA duplexes through interactions that involve the coiled-coil domain and the C-terminal IDR. Models of Mer2-Spp1-DNA complexes showed similar configurations with Mer2 and Spp1 residues binding the DNA backbone. Although these structural models require experimental validation, they are fully consistent with current data. First, based on AFM imaging and mutagenesis of IHO1 and Rec15 (Mer2 orthologs in mice and *S. pombe*), we previously proposed that the Mer2 coiled coil bridges DNA duplexes through electrostatic interactions along its length (Daccache et al. 2023), in agreement with the AlphaFold models. Second, the models are consistent with the known function of the Mer2-KRRR motif in promoting Mer2 DNA binding, condensation and DSB formation (Claeys Bouuaert et al. 2021). Third, the AlphaFold models identified the Spp1-R5K motif as putative DNA-binding residues, which we validated *in vitro*.

A DNA-binding activity of Spp1 was not anticipated based on its proposed loop-axis tethering function (Acquaviva et al. 2013; Sommermeyer et al. 2013). However, it has previously been noticed that Spp1 binds DNA *in vitro*, and this activity was mapped to a fragment of Spp1 containing residues 170-255 (Murton et al. 2010). Here, we identified the residues responsible for this activity.

In our hands, full-length Spp1 does not bind DNA by itself, but the C-terminal part of Spp1 alone does, suggesting that the N-terminus of Spp1 occludes the C-terminal DNA-binding domain. However, how this auto-inhibition may take place is not clear. Regardless, since mutation of the R5K motif compromises the assembly of Mer2-Spp1-DNA complexes *in vitro*, full-length Spp1 binds DNA in the context of its complex with Mer2, even if it does not interact with DNA by itself.

### An occlusion-compensation mechanism for Mer2-Spp1-DNA interactions

Based on truncation analyses, the C-terminal part of the Mer2 coiled coil had previously been implicated in binding Spp1 (Acquaviva et al. 2013) and DNA (Claeys Bouuaert et al. 2021). We therefore wondered how the recruitment of Spp1 and DNA might affect each other. Three general scenarios can be envisioned. Spp1 and DNA could bind independently to Mer2, they could compete for access to Mer2, or they could bind cooperatively (**Supplementary Figure 12**). First, independent and competition models are not compatible with a DNA-binding activity of Spp1. Second, titration experiments revealed no evidence for competition between Spp1 and DNA for Mer2 binding. In contrast, the presence of Spp1 slightly stimulated DNA binding and condensation, supporting a weak cooperativity model. The AlphaFold models fit with these observations, since Mer2 coiled coils can simultaneously accommodate Spp1 and DNA without major steric clashes, and interaction between Spp1 and DNA can account for the observed cooperativity. The cooperativity could be either reciprocal or unilateral (one-sided). The structural model fits best with a one-sided cooperativity model, where the presence of Spp1 facilitates DNA binding rather than *vice versa* (**Supplementary Figure 12E**).

In addition to independent, competition, and cooperativity models, another scenario can be considered, encompassing both negative and positive effects of Spp1 on DNA binding. We refer to this as an ‘occlusion-compensation’ model (**Supplementary Figure 12G**). Although the AlphaFold models suggest a harmonious coexistence of Spp1 and DNA around the Mer2 tetramers, a comparison of Mer2-Spp1 and Mer2-DNA models suggests that Spp1 occludes putative DNA-binding residues found along the Mer2 coiled coil. This suggests a model where Spp1 recruitment reduces the DNA-binding activity of Mer2, but this is compensated by Spp1-DNA interactions. Our data indicates that this is likely to be the case. Indeed, under the one-sided (Spp1 −> DNA) cooperativity model, mutation of the Spp1-R5K motif would be expected to reduce the DNA-binding activity of the complex but would not be expected to affect the ratio between Mer2-Spp1 and Mer2 nucleoprotein complexes. Instead, we found that mutating the Spp1-R5K motif does not strongly decrease DNA binding, but rather favors the formation of Mer2-DNA complexes at the expense of Mer2-Spp1-DNA complexes.

Hence, our interpretation is that the cost associated with Spp1 binding is compensated by interactions between the Spp1-R5K motif and DNA, such that the binding equilibrium is driven towards the formation of tripartite Mer2-Spp1-DNA complexes (**Figure 8A**). When the R5K motif is mutated, Spp1 recruitment leads to a net decrease in DNA binding, driving the equilibrium towards Mer2-DNA complexes. Consistent with this model, mutation of the Spp1-R5K motif drives the exclusion of Spp1 from Mer2 condensates *in vitro*.

Nevertheless, since mutating the R5K motif does not recapitulate the meiotic phenotype of a *spp1Δ* strain, the DNA-binding activity of Spp1 is not important for its meiotic function. Assuming that the R5K mutation also leads to a reduction of Spp1 binding to Mer2 condensates, as is the case *in vitro*, the implication is that the amount of Spp1 recruitment to Mer2 condensates is not a limiting factor in shaping the DSB landscape.

In summary, our work reveals new molecular insights into the role of Mer2 and Spp1 in the assembly of the meiotic DSB machinery.

## Materials and Methods

### Preparation of expression vectors

Plasmids are listed in **Supplementary Table 3**, and oligos are listed in **Supplementary Table 4**. The vectors used to express ^HisSUMO^Mer2 (pCCB750) and ^HisSUMO^eGFP-Mer2 (pCCB777) from *E. coli* were previously described (Claeys Bouuaert et al. 2021). The expression vector for ^His^Mer2 was generated by PCR amplification of the *MER2* coding sequence from pCCB750 with primers km001F and km001R and cloned into pET28b by Gibson assembly to yield pKM001. Expression vectors for Mer2 truncations were obtained by inverse PCR and self-ligation using template pCCB750 and primers cb1346 and cb1497 or cb1342 and dd121 to generate pDD057 (^HisSUMO^Mer2^41-314^) and pDD060 (^HisSUMO^Mer2^1-224^), respectively. Plasmid pDD057 was further used as template with primers cb1342 and dd121 to generate pDD065 (^HisSUMO^Mer2^41-224^).

The sequence of *SPP1* was amplified from yeast genomic DNA using primers pl001 and pl002 and cloned into pSMT3 vector by Gibson assembly to yield pPL001 (^HisSUMO^Spp1). *SPP1* with a Tev cleavage site was amplified using primers pl029 and pl030 and cloned by Gibson assembly in pMAL-c2x to yield pPL012 (^MBP^Spp1). *GST-Tev* was amplified from pCCB407 using primers pl035 and pl036 and cloned pEtDuet-1 to yield pPL015. The *SPP1* sequence with a Tev site was then amplified from pPL012 with primers pl037 and pl038 and cloned into pPL015 to yield pPL016 (^GST^Spp1). The expression vector for ^GST-His^SPP1 was created by amplifying the coding sequence of *SPP1* with primers km004F and pl038 and cloning by Gibson assembly into a XhoI and KpnI digested product of pPL016, yielding pKM004.

Sequences coding for ^HisSUMO^Mer2 and ^MBP^Spp1 were amplified from pCCB750 and pPL012 with primers pl031 and pl032, and pl033 and pl034 and cloned by Gibson assembly into pETduet to yield pPL014.

Expression vector for wild type ^MBP^Mer2 was obtained by amplification of *MER2* from pCCB750 with primers pl027 and pl028 and cloning into pMAL-c2x by Gibson assembly to yield pPL011. The expression vector for the Mer2-V195D mutant was obtained using pPL011 as a template by inverse PCR and self-ligation using pl053 and pl054 to yield pPL036.

Expression vector for Spp1^161-353^ truncation was generated by inverse PCR of pPL016 with primers pl145 and cb1486 and self-ligation to yield pPL094 (^GST^Spp1^161-353^). Expression vectors for full-length and truncated Spp1-R5K mutants were obtained using pPL016 and pPL094 as inverse PCR templates using primers pl146 and pl147 and self-ligation to yield respectively pPL099 (^GST^Spp1-R5K) and pPL095 (^GST^Spp1^161-353^-R5K).

Co-expression vectors for ^HisSUMO^Mer2^138-224^ and ^MBP^Spp1^161-353^ were made as follows. First, the sequences coding for the Mer2 and Spp1 fragments were amplified from pCCB750 and pPL012 using primers pl078 and pl079, and pl080 and pl081 and cloned into pSMT3 and pMAL-c2x by Gibson assembly to yield pPL041 and pPL042. Then the sequences coding for ^HisSUMO^Mer2^138-224^ and ^MBP^Spp1^161-353^ were amplified from pPL041 and pPL042 using primers pl082 and pl083, and pl104 and pl105 and cloned separately into pEtDuet-1 by Gibson assembly to yield pPL045 and pPL053. Finally, the co-expression vector for ^HisSUMO^Mer2^138-224^ and ^MBP^Spp1^161-353^ truncations was obtained by restriction digestion of pPL045 and pPL053 using XhoI and HindIII and ligation of the fragments of interest to yield pPL054.

Point mutations for the analysis of the Mer2-Spp1 interaction surface were obtained by inverse-PCR and self-ligation using pPL054 (^HisSUMO^Mer2^138-224, MBP^Spp1^161-353^) as a template. PCR primers and resulting plasmids for each mutant are as follows: Spp1-Q317A (pl112 and pl113, pPL064), Spp1-Q318A (pl114 and pl115, pPL065), Spp1-S324A (pl134 and pl135, pPL066), Spp1-R328A (pl132 and pl133, pPL067), Spp1-L332A (pl118 and pl119, pPL068), Spp1-R336A (pl120 and pl121, pPL069), Spp1-Q339A (pl122 and pl123, pPL070), Spp1-I342A (pl124 and pl125, pPL071), Mer2-N184A (pl140 and pl141, pPL072), Mer2-Q188A (pl138 and pl139, pPL073), Mer2-E190A (pl136 and pl137, pPL074), Mer2-V195D (pl053 and pl054, pPL075), Mer2-D202A (pl116 and pl117, pPL076), Mer2-T208A (pl128 and pl129, pPL077), Mer2-R209A (pl126 and pl127, pPL078), Mer2-E214A (pl130 and pl131, pPL079), and Mer2-V195A (pl053 and pl144, pPL089).

The co-expression vector for Spp1^307-353^ and Mer2^138-224^ truncations was obtained by inverse PCR and self-ligation using pPL054 as a template and primers pl143 and cb1486 to yield pPL090.

The expression vector for Spp1 fused with mScarlet was obtained by amplifying mScarlet sequence from pCCB785 (Claeys Bouuaert et al. 2021) with primers pl057 and pl058 and cloned in pPL016 by Gibson assembly to yield pPL032. Expression vector for wild-type Mer2 fused to eGFP was previously described (Claeys Bouuaert et al. 2021) and V195D mutant was obtained using this construct (pCCB777) as a template by inverse PCR and self-ligation using pl053 and pl054 to yield pPL037.

### Expression and purification of recombinant proteins

For the expression of Spp1 proteins (GST-tagged Spp1 (WT and R5K), untagged Spp1 (full-length and 161-353 truncations, WT and 5KR mutants), His-tagged Spp1, mScarlet-Spp1 (WT and R5K), vectors were transformed in *E. coli* C41 cells and plated on LB plates containing the appropriate antibiotic. Cells were then cultured at 37°C to an optical density (OD_600_) of 0.6. All the proteins were expressed as a fusion with an N-terminal GST tag cleavable with TEV protease. Expression was carried out at 18°C overnight with 0.25 mM isopropyl β-D-1-thiogalactopyranoside (IPTG). Cell pellets were frozen in liquid nitrogen and kept at −80°C until use. All the purification steps were carried out at 0-4°C. Cells were then resuspended in lysis buffer (50 mM HEPES pH7.5, 300 mM NaCl, 5% glycerol, 0.1% Triton, 5 mM beta-mercaptoethanol, 1 mM PMSF), lysed by sonication and centrifuged at 43,000 *g* for 30 minutes. The cleared extract was loaded onto 1 mL pre-equilibrated Glutathione sepharose resin (Cytiva). The column was washed extensively with lysis buffer. Proteins that retained the GST tag were then eluted with 5 mL lysis buffer containing 10 mM reduced Glutathione. For the versions without GST, the tag was cleaved on the column by overnight treatment with Tev protease. The flow-through was then loaded on a Superdex 200 Increase 10/300 GL column preequilibrated with gel filtration buffer (20 mM HEPES pH 7.5, 500 mM NaCl, 10 % glycerol, 0.1 mM DTT). For all purifications, after gel filtration, fractions containing the protein were concentrated using 10-kDa cutoff Amicon centrifugal filters (Millipore). Aliquots were flash frozen in liquid nitrogen and stored at −80°C.

For the expression of Mer2 proteins (untagged Mer2 (WT and V195D), Alexa-labeled Mer2, truncations (Mer2^138-224^ and Mer2^1-224^), eGFP-tagged Mer2 (WT and V195D)) and for co-expression of Mer2-Spp1 complexes, vectors were transformed in *E. coli* BL21 DE3 cells and plated on LB plates containing the appropriate antibiotic. Cells were then cultured in liquid medium at 37°C to OD_600_ of 0.6. All constructs expressed Mer2 with an N-terminal 6His-SUMO tag, cleavable with Ulp1. Co-expression vectors also expressed Spp1 with an N-terminal MBP tag, cleavable by TEV protease. Expression was carried out at 30°C for 3 hours with 1 mM IPTG for Mer2 and at 18°C overnight with 0.25 mM IPTG for the complexes. Cell pellets were frozen in liquid nitrogen and kept at −80°C until use. All the purification steps were carried out at 0-4°C. Cells were resuspended in nickel buffer (25 mM HEPES-NaOH pH 7.5, 1 M NaCl, 10% glycerol, 0.1 mM DTT, 40 mM imidazole, 1 mM PMSF), lysed by sonication and centrifuged at 43,000 *g* for 30 minutes. The cleared extract was loaded onto 1 mL pre-equilibrated Ni-NTA resin (Thermo Scientific). The column was washed extensively with nickel buffer then eluted in buffer containing 250 mM imidazole. To remove the His-SUMO tag of Mer2, complexes were incubated for 3 hours with Ulp1. For Mer2 protein labeled with Alexa Fluor 488 (Invitrogen no. A10235), the Ulp1-treated protein was dialysed overnight in nickel buffer without imidazole, followed by fluorophore conjugation at room temperature for 1 hour before gel filtration. Alexa Fluor 488 has a tetrafluorophenyl ester moiety that reacts with primary amines. All the proteins were then loaded either on a Superdex 200 Increase 10/300 GL column (Mer2) or on a Superose 6 Increase 10/300 GL column (Mer2-Spp1 complexes) preequilibrated with gel filtration buffer (25 mM HEPES-NaOH pH 7.5, 300 mM NaCl, 10% glycerol, 1 mM DTT, 5 mM EDTA). For all purifications, after gel filtration, fractions containing the protein were concentrated using 10-kDa cutoff Amicon centrifugal filters (Millipore). Aliquots were flash frozen in liquid nitrogen and stored at −80°C.

For SEC-SAXS analyses, ^HisSUMO^Mer2 - ^MBP^Spp1, Mer2 - ^MBP^Spp1, and Mer2^138-224^ - ^MBP^Spp1^161-353^ complexes were obtained by co-transformation of the appropriate expression vector, affinity chromatography with Ni-NTA resin, followed by size-exclusion chromatography using SAXS buffer (25 mM HEPES-NaOH pH 7.5, 500 mM NaCl, 5 mM EDTA, 5 % glycerol, 1 mM TCEP). Mer2^41-224^-Spp1 and Mer2^41-224^-Spp1^161-353^ complexes were obtained by mixing purified Mer2 and Spp1 subunits at a 4:6 ratio, followed by size exclusion chromatography with SAXS buffer.

### Pulldown assays

For Ni-NTA pulldowns made from cell lysates, tagged Mer2 and Spp1 truncations were co-expressed in 50 mL of *E. coli* BL21 cultures and purified by affinity chromatography on nickel resin following a procedure similar to that described above. Cells were lysed by sonication and centrifuged at 15,000 rpm for 15 minutes. The cleared extract was loaded onto 100 μL of pre-equilibrated Ni-NTA resin (Thermo Scientific) in nickel buffer containing 25 mM HEPES-NaOH pH 7.5, 1 M NaCl, 0.1% Triton X-100, 10% glycerol, 0.1 mM DTT, 40 mM imidazole. After 30 minutes incubation on a rotating wheel at 4°C, the resin was loaded on a column, washed with 15 mL nickel buffer and then eluted in buffer containing 250 mM imidazole. Eluted proteins were boiled in 1× Laemmli buffer and analyzed by SDS–PAGE.

GST pulldowns were made with the indicated concentrations of purified proteins in 40 μL reactions in buffer containing 20 mM HEPES-NaOH pH 7.5, 300 mM NaCl, 5% glycerol, 1 mM DTT. After a 2 hours incubation on ice, the proteins were mixed with 100 μL pre-equilibrated Glutathione sepharose resin (Cytiva) and left for 30 minutes on ice with occasional mixing. The resin was then centrifuged at 1500 g for 1 minute and washed three times with 1 mL pulldown buffer. Both input and output were resuspended in the same volume of Laemmli buffer and analyzed by SDS-PAGE.

### SEC-MALS

Light scattering data were collected using a Superose 6 increase 10/300 GL Size Exclusion Chromatography (SEC) column connected to an AKTA Pure Chromatography System (Cytiva). The elution from SEC was monitored by a differential refractometer (Optilab, Wyatt), and a static and dynamic, multi-angle laser light scattering (LS) detector (miniDAWN, Wyatt). The SEC–UV/LS/RI system was equilibrated in 25 mM HEPES-NaOH pH 7.5, 300 mM NaCl, 1 mM DTT, 5 mM EDTA at a flow rate of 0.4 mL/min. The weight average molecular masses were determined across the entire elution profile at intervals of 0.5 seconds from static LS measurement using ASTRA software.

### Functionalization of AFM tips

AFM experiments were performed using Pyrex-nitride probe with a triangular gold-coated pyramidal silicon nitride PNP-TR-AU50 cantilever (Nanoworld) calibrated using the thermal noise method. For tip functionalization, the cantilevers were rinsed with ethanol, dried under a gentle nitrogen stream and cleaned for 15 minutes by UV and ozone treatment (Jetlight). After that, tips were incubated overnight in a polyethylene glycol (PEG) solution prepared by mixing 0.05 mM HS-(CH_2_)_11_-EG_3_-OH with 0.05 mM HS-(CH_2_)_11_-EG_3_-NTA dissolved in ethanol in a 99:1 (v/v) ratio. The tips were then rinsed with ethanol, dried with nitrogen, and treated with 50 mM NiCl_2_ for 30 minutes (Pfreundschuh et al. 2015).

The tips were functionalized with Spp1 protein through an N-terminal His6 tag. The cantilevers were placed on Parafilm (Bemis NA), incubated with 40 μL protein solution (0.1 mg/mL in buffer containing 20 mM HEPES-NaOH pH 7.5, 300 mM NaCl, 10% glycerol, 1 mM DTT and 5 mM EDTA) for 1 hour, and washed three times with protein binding buffer (20 mM Tris-HCl, 10 mM MgCl_2_, and 100 mM NaCl).

### Preparation of protein coated model surfaces

Proteins were immobilized on gold-coated silicon surfaces (Platypus) using NHS-EDC chemistry (Koehler et al. 2021). The surfaces were first rinsed with ethanol, dried under nitrogen, and cleaned by UV-ozone treatment for 15 minutes. The surfaces were then incubated overnight in a solution containing alkanethiols (90% 1 mM 11-mercapto-1-undecanol and 10% 1 mM 16-mercaptohexadecanoic acid in ethanol). Next, the surfaces were rinsed with ethanol, dried under a gentle nitrogen stream, and immersed in a solution containing 25 mg/mL dimethylaminopropyl carbodiimide (EDC) and 10 mg/mL N-hydroxy succinimide (NHS) dissolved in H_2_O. Finally, the chemically activated surfaces were incubated with 50 µL of the respective protein (15 µg/mL in buffer containing 20 mM HEPES-NaOH pH 7.5, 300 mM NaCl, 10% glycerol, 1 mM DTT and 5 mM EDTA) on Parafilm for 1 hour at 4 °C, washed with protein binding buffer, and used on the same day.

### Force-distance curve-based AFM

The acquisition of force-distance curves by AFM on model surfaces was carried out in protein binding buffer at room temperature, using the tips and surfaces described above. AFM tips were vertically approached and retracted from the surface while recording the force. A Multimode Nanoscope VIII (Bruker) was used. Areas of 25 μm^2^ (1024 pixels/map) were scanned, the tip approach and retraction speeds were fixed at 1 μm/s, the ramp size was set to 500 nm, a setpoint force of 500 pN, and a resolution of 32 × 32 pixels. Curves were assessed for specific adhesion events individually, and the proportion of curves showing binding versus those that did not was used to calculate the binding frequency.

Dynamic force spectroscopy (DFS) experiments on model surfaces were conducted using a ForceRobot 300 (Bruker) under the same conditions as the binding frequency measurements, with a varying retraction velocity of 0.1, 0.2, 1, 5, 10, and 20 μm/s. Force curves were analyzed using JPK Data Processing software (version 6.1.149). The results were plotted as DFS graphs and rupture force histograms for distinct loading rates ranges using Origin software (OriginLab), and the data were fitted to the Bell-Evans models as described elsewhere to extract the kinetic off rate (Rankl et al. 2011; Alsteens et al. 2017). Briefly, the relationship between rupture force and loading rate is described by the following equation:

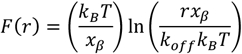

where *F(r)* is the most probable rupture force, *k*_*B*_ Boltzmann’s constant, *T* the temperature in Kelvin, *x*_*β*_ the distance to the transition state, *r* the loading rate and *k*_*off*_ the off-rate constant at zero force.

For kinetic on-rate (k_on_) analysis, the binding frequency was measured by varying the contact time (t) (the time the AFM tip is in contact with the surface). Briefly, the relationship between interaction time (τ) and binding frequency (BF) is described by the following equation:

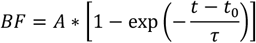

where *A* represents the maximum BF and *t*_*0*_ the lag time. Data fitting and τ extraction were performed using Origin software. The *k*_*on*_ value was subsequently calculated using the following equation:

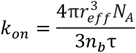

where, *r*_*eff*_ is the effective radius of the sphere, *n*_*b*_ is the number of binding partners, and *N*_*A*_ is the Avogadro constant. The effective volume is defined as *V*_*eff*_ = *(4πr*_*eff*_^*3*^*)/3*, representing the volume in which the interaction can take place. The parameters used for *k*_*on*_ calculation were as follows: *n = 1* molecule of Spp1; τ = 75±12 ms, extracted from the curve fit (**Figure 1D**); *r*_*eff*_ = 14.5 nm, calculated based on the AlphaFold model of Spp1 that yields a distance between the N-terminal His tag and C-terminal Mer2 binding site of about 11.5 nm, to which we added 3 nm for the PEG linker. Quantitative parameters for *k*_*on*_ and *k*_*off*_ are presented in **Supplementary Table 1**.

### SEC-SAXS

All experiments were performed at the SOLEIL BioSAXS beamline SWING (Gif-sur-Yvette, France). SEC-SAXS data were collected in HPLC mode using a Shodex KW404-4F column pre-equilibrated with SAXS buffer (25 mM HEPES-NaOH pH 7.5, 500 mM NaCl, 5 mM EDTA, 5% glycerol, 1 mM TCEP). Samples were concentrated on-site to ~3-5 mg/mL using 10-kDa cutoff centrifugal filters (Amicon). Ninety-microliter samples were injected and eluted at a flow rate of 0.2 mL/min while scattering data was collected with an exposure time of 990 msec and a dead time of 10 msec. The scattering of pure water was used to calibrate the intensity to absolute units. Data reduction was performed using FoxTrot. Data were processed using BioXTas RAW (Hopkins et al. 2017) and analyzed using RAW and the ATSAS package (Manalastas-Cantos et al. 2021). The information on data collection and derived structural parameters is summarized in **Supplementary Table 2**. *Ab initio* calculations were performed with the DENSS suite (Grant 2018) and P2 symmetry was imposed during shape reconstruction.

The molecular models of the various Mer2-Spp1 complexes were generated from the AlphaFold models obtained in this work and refined in Xplor-NIH v 2.49 (Schwieters et al. 2003). In all refinement runs, the positions of the Mer2 coiled-coil domain and the interacting Spp1 helices 1 and 2 were kept fixed; SUMO globular domains treated as rigid body groups, while the intervening linkers and terminal tails given full torsional degrees of freedom. The computational protocol comprised an initial simulated annealing step followed by the side-chain energy minimization. The total minimized energy function consisted of the standard geometric (bonds, angles, dihedrals, and impropers) and steric (van der Waals) terms, a knowledge-based dihedral angle potential (Schwieters et al. 2003), and a SAXS energy term incorporating the experimental data (Schwieters and Clore 2014). In each refinement run, 100 structures were calculated, and 10 lowest-energy solutions retained for further analysis. The agreement between the experimental and calculated SAXS curves (obtained with the calcSAXS helper program, which is part of the Xplor-NIH package) was assessed by calculating the χ2:

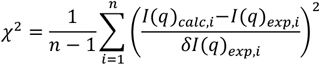

where *I*(*q*)_*calc,i*_ and *I*(*q*)_*exp,i*_ are the scattering intensities at a given q for the calculated and experimental SAXS curves, *δI*(*q*)_*exp,i*_ is an experimental error on the corresponding *I*(*q*)_*exp,i*_ value, and n is the number of data points defining the experimental SAXS curve. The models were fitted into the ab initio densities with UCSF ChimeraX (Meng et al. 2023).

### DNA substrates and gel shift assays

A fluorescent substrate that mimics a Holliday Junction was generated by annealing complementary oligos (cb923, cb924, cb925, cb934). Oligos were mixed in equimolar concentrations (10 μM) in STE (100 mM NaCl, 10 mM Tris-HCl pH 8, 1 mM EDTA), heated and slowly cooled on a PCR thermocycler (98°C for 3 min, 75°C for 1 h, 65°C for 1 h, 37°C for 30 min, 25°C for 10 min).

Binding reactions (20 μl) were carried out in 25 mM Tris-HCl pH 7.5, 10% glycerol, 100 mM NaCl, 2 mM DTT, and 1 mg/mL BSA. Reactions contained 10 nM fluorescently labeled substrate and the indicated concentration of protein. Complexes were assembled for 30 minutes at 30°C and separated by gel electrophoresis. Binding reactions were separated on 5% TAE-polyacrylamide gels at 150 V for 2.5 hours, and gels were visualized using a Typhoon scanner (Cytiva).

### In vitro condensation assays

Condensation was induced by threefold dilution of Spp1 and Mer2 proteins in buffer containing DNA and no salt to reach final 20 μL reactions that contained 3 nM (100 ng) supercoiled plasmid DNA (pUC19), 25 mM Tris-HCl (pH 7.5), 10% glycerol, 100 mM NaCl, 2 mM DTT, 1 mg/mL BSA, and 5% PEG 8000. After 30 minutes at 30°C with occasional mixing, 4 μL was dropped onto a microscope slide and covered with a coverslip. Images were captured on a Zeiss Axio Observer with a 100×/1.4 NA oil immersion objective. Images were analyzed with ImageJ using a custom-made script. In brief, a fixed threshold was applied to 129.24 × 129.24-μm (2048 × 2048-pixel) images. The intensity inside the focus mask was integrated. Data points represent averages of 10 images per sample. Data were analyzed using GraphPad Prism 9.

### Yeast strain construction

Yeast strains were from the SK1 background. All strains used in this study are listed in **Supplementary Table 5**. The *SPP1-R5K* mutant was constructed by transformation of a DNA fragment obtained by overlap extension PCR. The 3’ end of the *SPP1* gene containing the R5K mutation was amplified from plasmid pPL099 with primers pl160 and pl161. The KanMX6 cassette was amplified with primers pl162 and pl163. The two PCR products were purified by gel extraction and combined in the overlap extension PCR using primers pl160 and pl163. The resulting product was transformed into CBY006 cells and transformants selected with G418. The *SPP1-R5K* mutation was confirmed by PCR and sequencing of the *SPP1* genomic region and strains carrying this allele were subsequently produced by mating and tetrad dissection.

### Southern blot analysis of DSBs

Meiotic DSB analysis by Southern blotting was performed as described (Murakami et al. 2009). Briefly, diploid strains were grown overnight at 30 °C on YPG plates (1% yeast extract, 2% peptone, 3% glycerol) and then streaked on YPD plates (1% yeast extract, 2% peptone, 2% dextrose). Two days later, diploid colonies were transferred to liquid YPD and cultured overnight with vigorous shaking. Saturated cultures were diluted in SPS (0.5% Yeast Extract, 1% peptone, 0.67% yeast nitrogen base, 1% potassium acetate, 50 mM potassium phtalate, and antifoam) to a density of 1.3×10^6^ cells/ml. After 7 hours at 30 °C, cultures reached a concentration of 1-1.3×10^7^ cells/ml. Cells were diluted again in SPS to a final density of 3.25×10^5^ cells/ml and incubated for 13-16 hours until they reached a density of 3×10^7^ to 4×10^7^ cells/ml. Meiosis was induced by transferring the cells to 2% KOAc, supplemented with amino acids and 0.01% polypropylene glycol, and cells were harvested at the indicated time. After DNA extraction, 1-1.2 µg of genomic DNA was digested either with HindIII-HF or KpnI-HF (NEB), for Southern blot analysis at *CYS3* and *PES4* hotspots, respectively, and separated on a 1% TBE-agarose SEA-KEM LE gel for 16 hours at 70 V. DNA was transferred to a positively charged nylon membrane (Hybond-N+ from Amersham) by vacuum transfer, hybridized with *CYS3* probe (amplified with primers: TTTGAAGGTCACCGACATCC and CGATTAGTTGGTGGCTTGTT) or with the *PES4* probe (amplified with primers AGCTTATCAAACTGTGTGCA and GGCTGGGGATCTTGAAAGTA). Films were exposed for 3 days and developed by autoradiography using a Typhoon Biomolecular Imager (Cytiva).

## End Matter

### Author Contributions and Notes

C.C.B designed the research; generated AlphaFold models and analyzed data; P.L. designed, executed, and analyzed all experiments except as noted; D.A.M. performed Southern blot experiments; V.G.G., S.M. and S.D. performed AFM experiments under the supervision of D.A.; K.M. performed preliminary AFM experiments and analyzed data with A.R; Y.G.-J.S. performed SAXS experiments with P.L. and analyzed data with A.N.V.; C.C.B wrote the paper with input from all authors. C.C.B. supervised the research and secured funding. The authors declare no competing interests.

## Acknowledgments

We thank Dima Daccache for help with protein purifications, and Laurent Acquaviva for the *spp1 Δ* strain. We thank Javier Perez and Aurélien Thureau of the SWING beam line at SOLEIL synchrotron for outstanding technical support. This work was supported by the European Research Council under the European Union’s Horizon 2020 research and innovation program (ERC grant agreement 802525 to CCB), and the Fonds National de la Recherche Scientifique (PDR grant T.0031.22 to CCB and PDR grant T.0003.23 to DA). KM is funded by an FNRS Aspirant fellowship (project FC42859). V.G.G. is a postdoctoral researcher at the FNRS. DA is a Senior Research Associate at the FNRS. CCB is a FNRS Research Associate.

## Supplementary Information

**Supplementary Figure 1:**
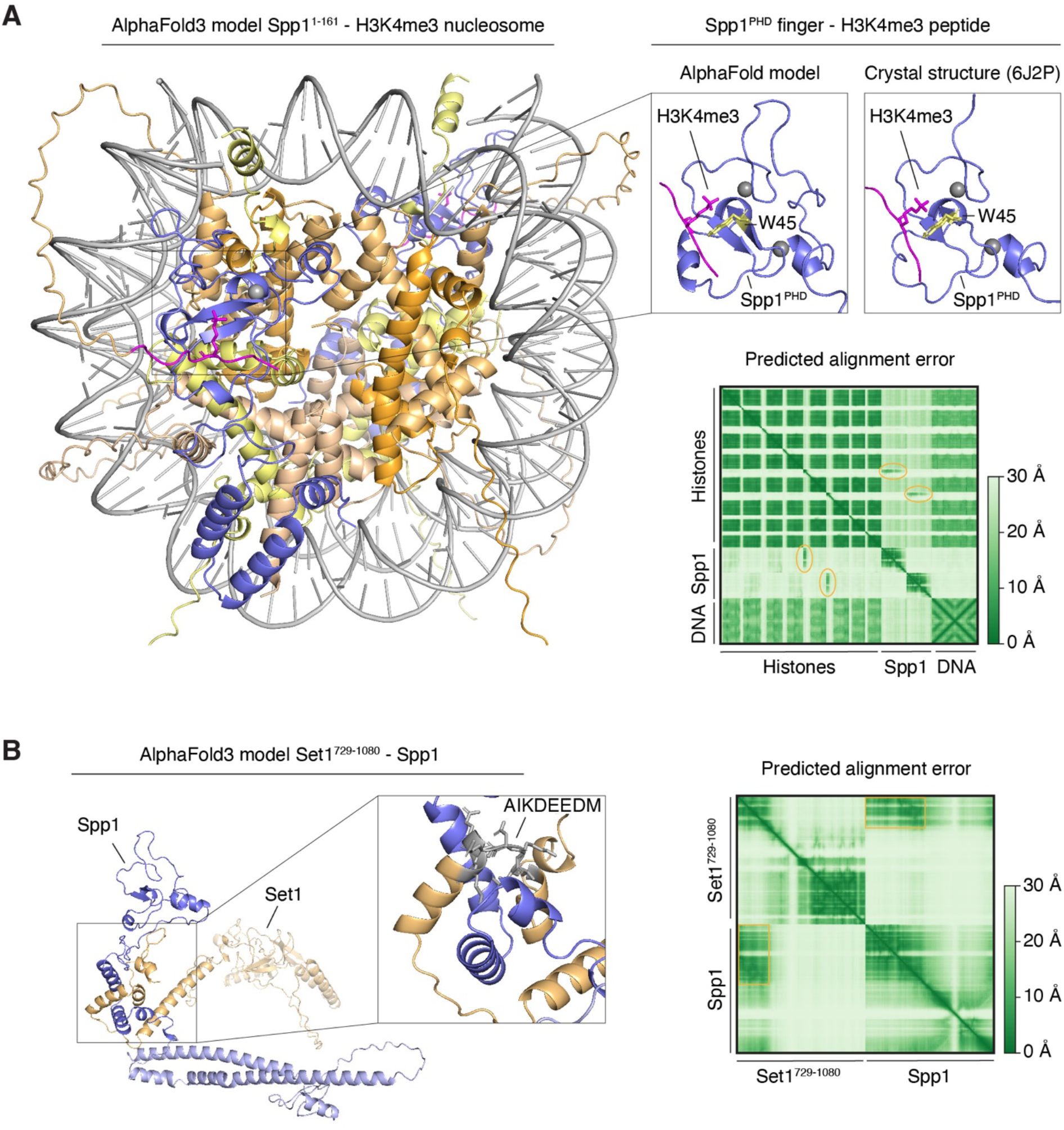
AlphaFold models of Spp1 bound to H3K4 trimethylated nucleosome and Set1. **(A)** AlphaFold3 model of Spp1^1-160^ bound to a H3K4me3 marked nucleosome. H2A (HTA1) is yellow, H2B (HTB1) is wheat, H3 (HHT1) is light orange, H4 (HHF1) is bright orange, DNA is grey, the N-terminal H3 tail is magenta, and Spp1 is blue. Top right: Comparison of the AlphaFold model and crystal structure (6J2P) (He et al. 2019) of the Spp1 PHD finger bound to a lysine 4 trimethylated H3 histone tail. Spp1 residue W45 previously shown to be essential for H3K4me3 binding (Shi et al. 2007; Sommermeyer et al. 2013) is shown in yellow. Zinc ions are shown as grey spheres. Bottom right: Predicted alignment error (PAE) plot with interaction between Spp1^PHD^ and H3K4me3 highlighted (orange ovals). **(B)** AlphaFold3 model of Spp1 (blue) bound to Set1 (yellow). Zoom: Set1 motif A_767_IKDEEDM_774_ important for Spp1 interaction (Adam et al. 2018) is in grey. Right: PAE plot with regions of interaction between Spp1 and Set1 highlighted (orange squares).

**Supplementary Figure 2:**
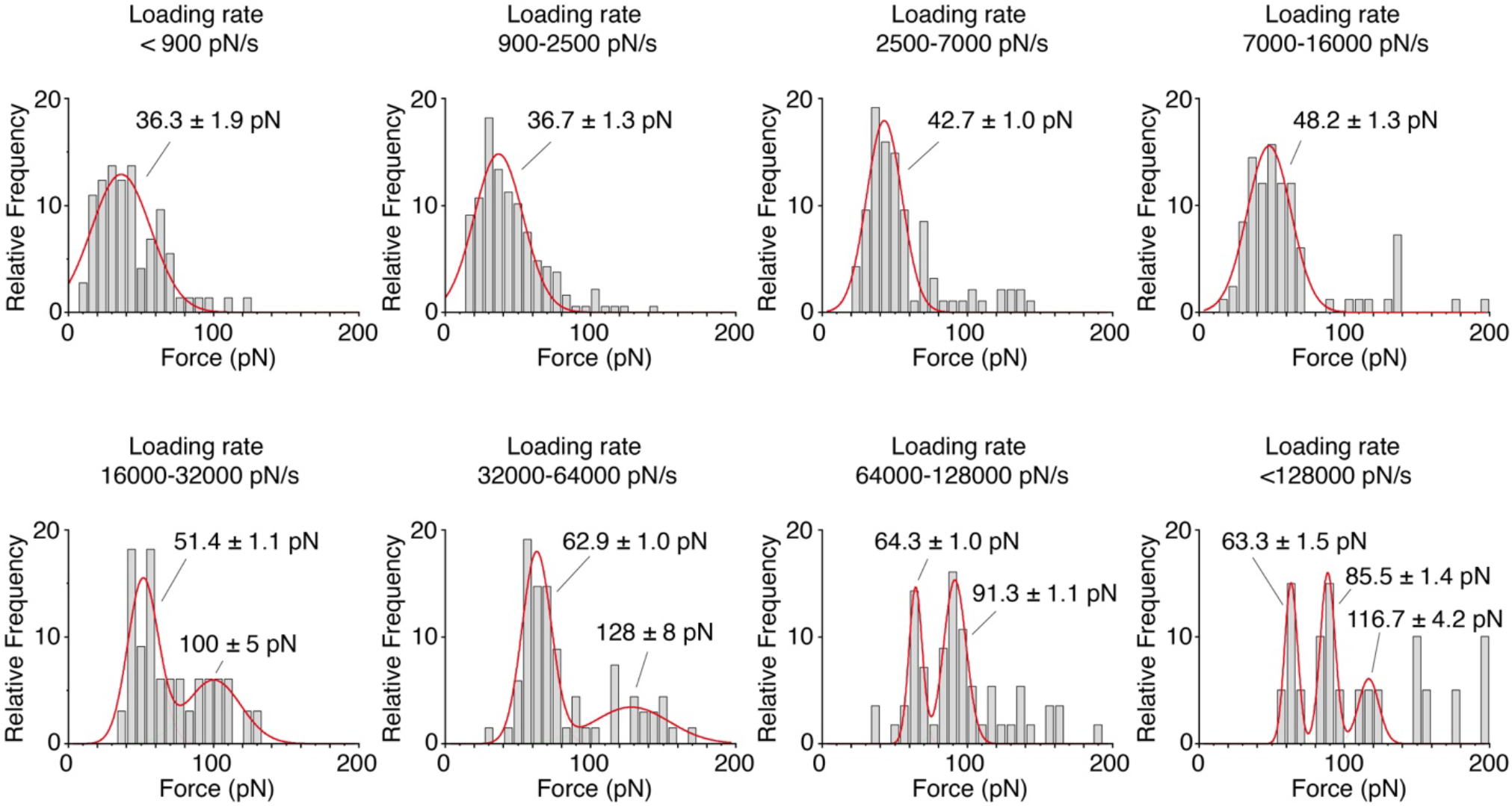
AFM rupture forces at different loading rates. Histograms of AFM rupture forces between Spp1 and Mer2 obtained at different ranges of loading rates. Solid lines represent multipeak Gaussian fits to the data. The rupture force increases as a function of loading rate. At loading rates above 10^4^ pN/s, additional peaks of higher rupture forces appear, probably caused by the disruption of the interaction between the histidine tag and Ni^2+^-NTA.

**Supplementary Figure 3:**
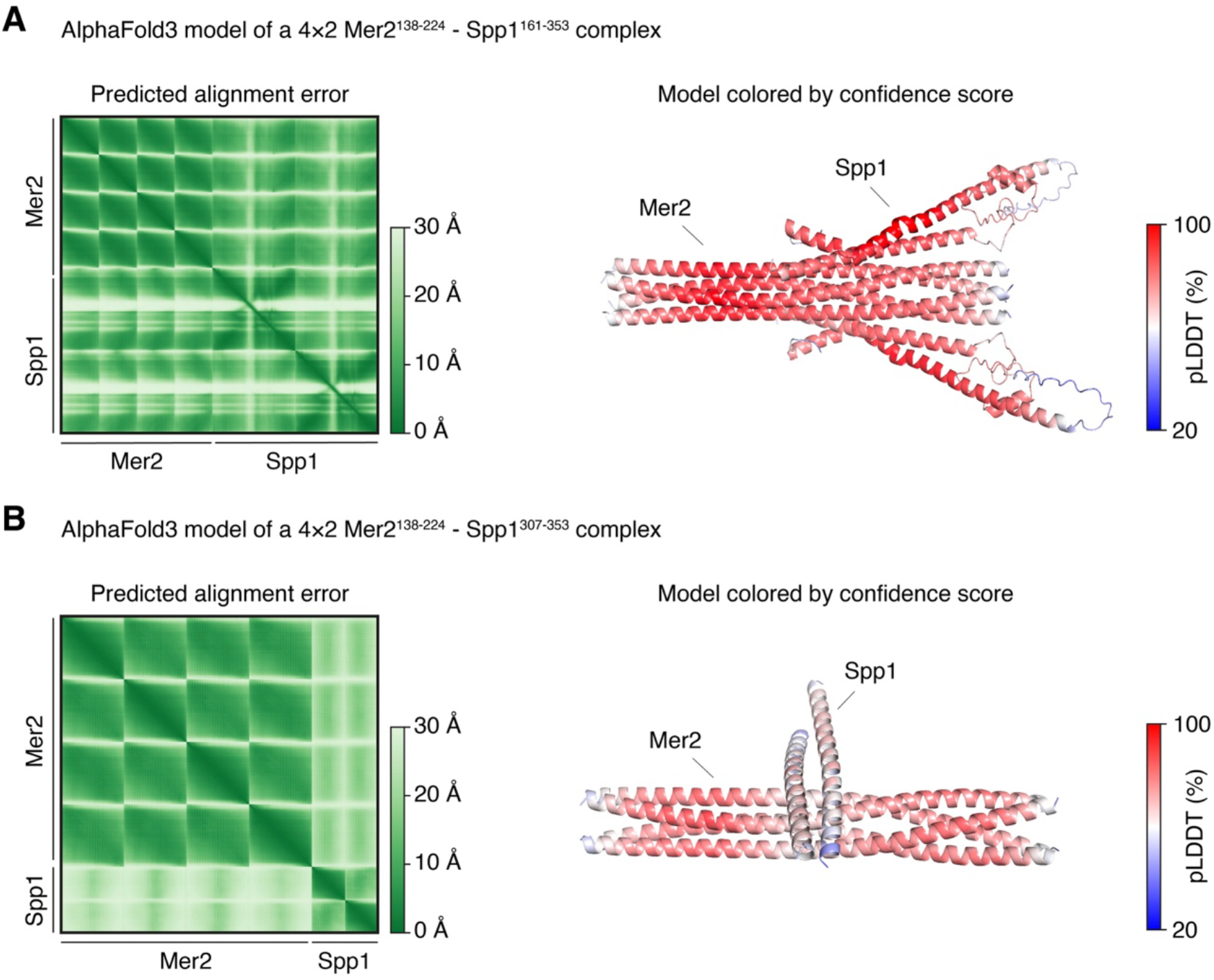
Quality assessment of the minimal Mer2-Spp1 AlphaFold models. AlphaFold3 models of 4×2 Mer2-Spp1 complexes including the indicated protein fragments. Left, predicted alignment error plot (low values indicate high confidence). Right, models colored by confidence score (red indicates high confidence). Input sequences are provided in **Supplementary Table 6**.

**Supplementary Figure 4:**
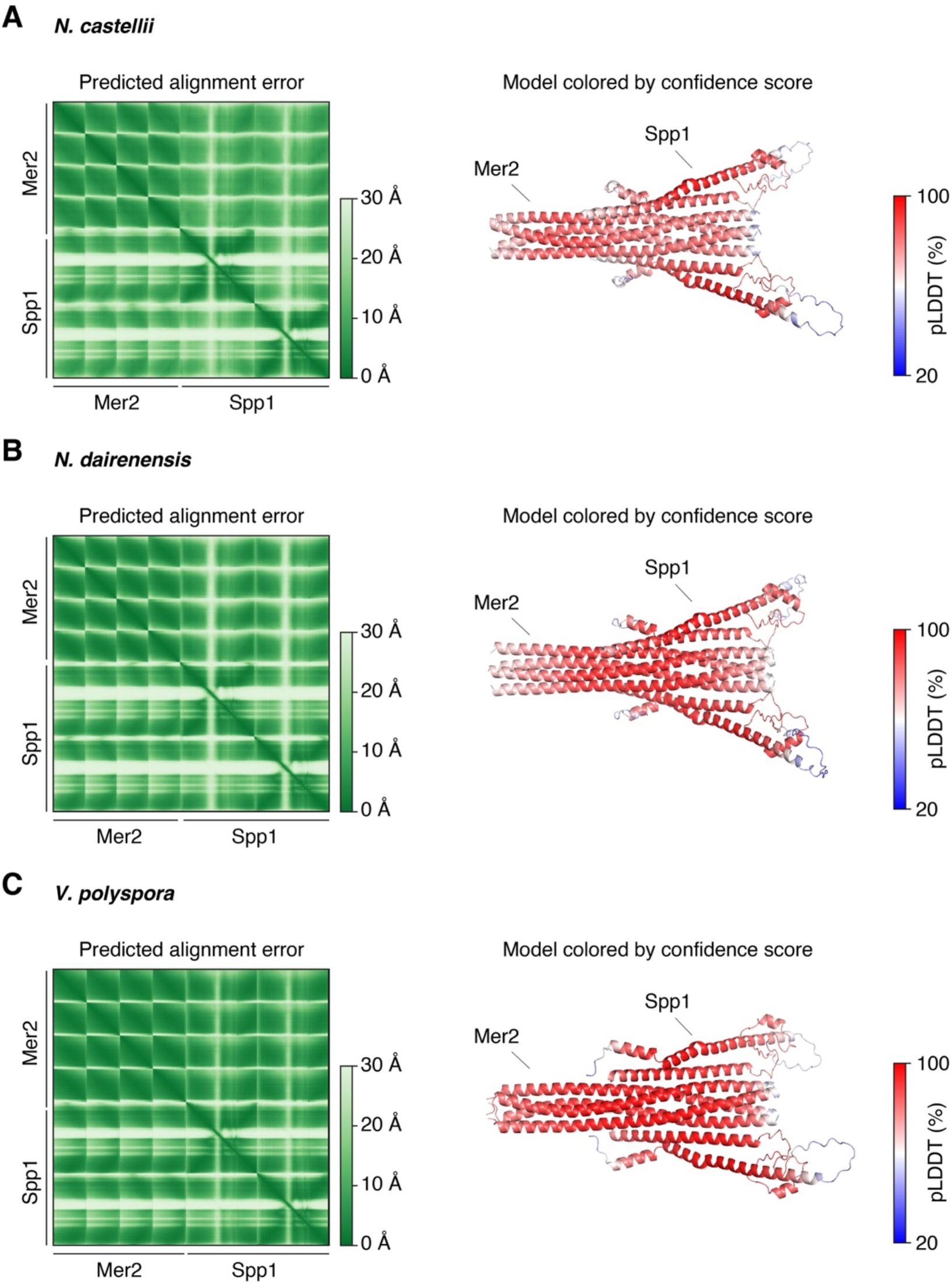
Structural prediction of the Mer2-Spp1 interface from different yeast species. AlphaFold3 models of the Mer2-Spp1 interaction domain from **(A)** *Naumovozyma castellii*, (XP_003676113.1, XP_003670290.1), **(B)** *Naumovozyma dairenensis* (XP_003669210.1, XP_003670290.1), **(C)** *Vanderwaltozyma polyspora* (XP_001647040.1, XP_001642735.1). Input sequences are provided in **Supplementary Table 6**. Left, predicted alignment error plot (low values indicate high confidence). Right, models colored by confidence score (red indicates high confidence).

**Supplementary Figure 5:**
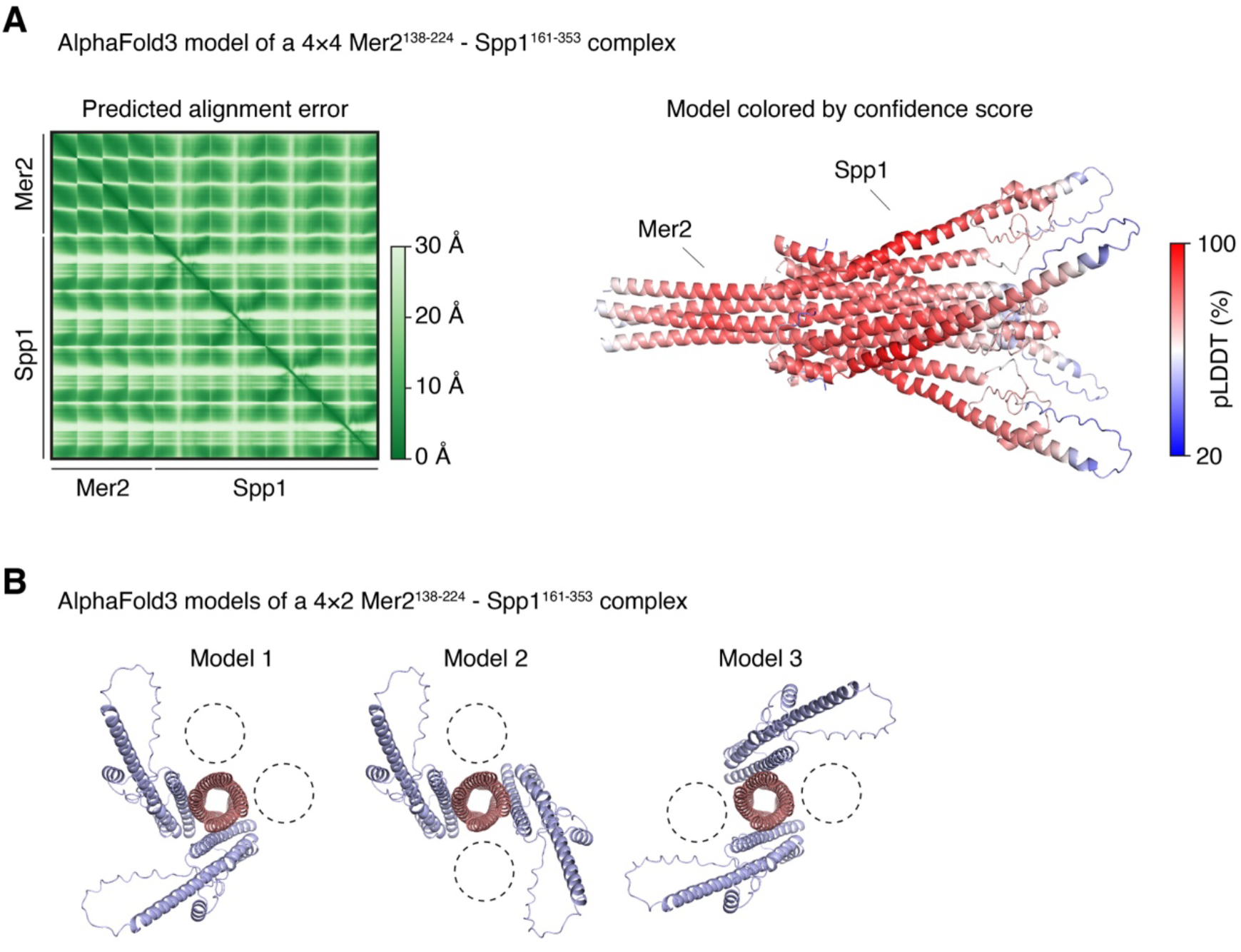
The AlphaFold model is consistent with a 4×4 Mer2-Spp1 complex. **(A)** AlphaFold3 model of the interaction domain of Mer2-Spp1 with a 4×4 stoichiometry. Left, predicted alignment error plot (low values indicate high confidence). Right, models colored by confidence score (red indicates high confidence). **(B)** Comparison of three 4×2 models, illustrating the arbitrary positioning of two Spp1 monomers (blue) around the Mer2 coiled coil (pink). Vacant binding sites are shown as dashed circles.

**Supplementary Figure 6:**
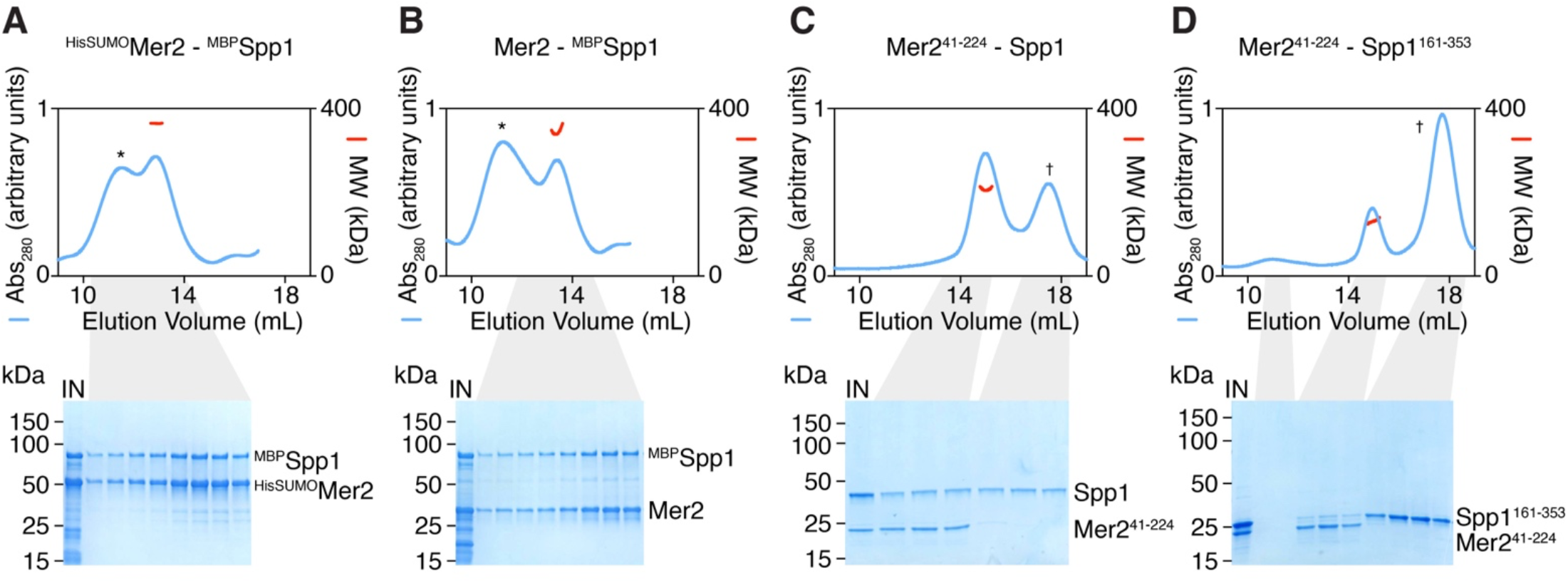
SEC-MALS analyses of Mer2-Spp1 complexes. SEC-MALS analyses (top) and SDS-PAGE analyses of SEC fractions (bottom) of tagged, untagged full-length and truncated Mer2-Spp1 complexes. Blue traces show the absorbance at 280 nm (arbitrary units) and red traces are molar mass measurements across the peak of interest. Samples A and B were produced by co-expression of tagged complexes in *E. coli* and purification by affinity chromatography. Samples C and D were produced by mixing purified truncated Mer2 with and excess of full length or truncated Spp1. Peaks labeled (*) correspond to higher-order oligomers, observed only when full-length Mer2 is present. Peaks labeled (†) correspond to an excess of Spp1. See summary of the data in **Figure 4A**.

**Supplementary Figure 7:**
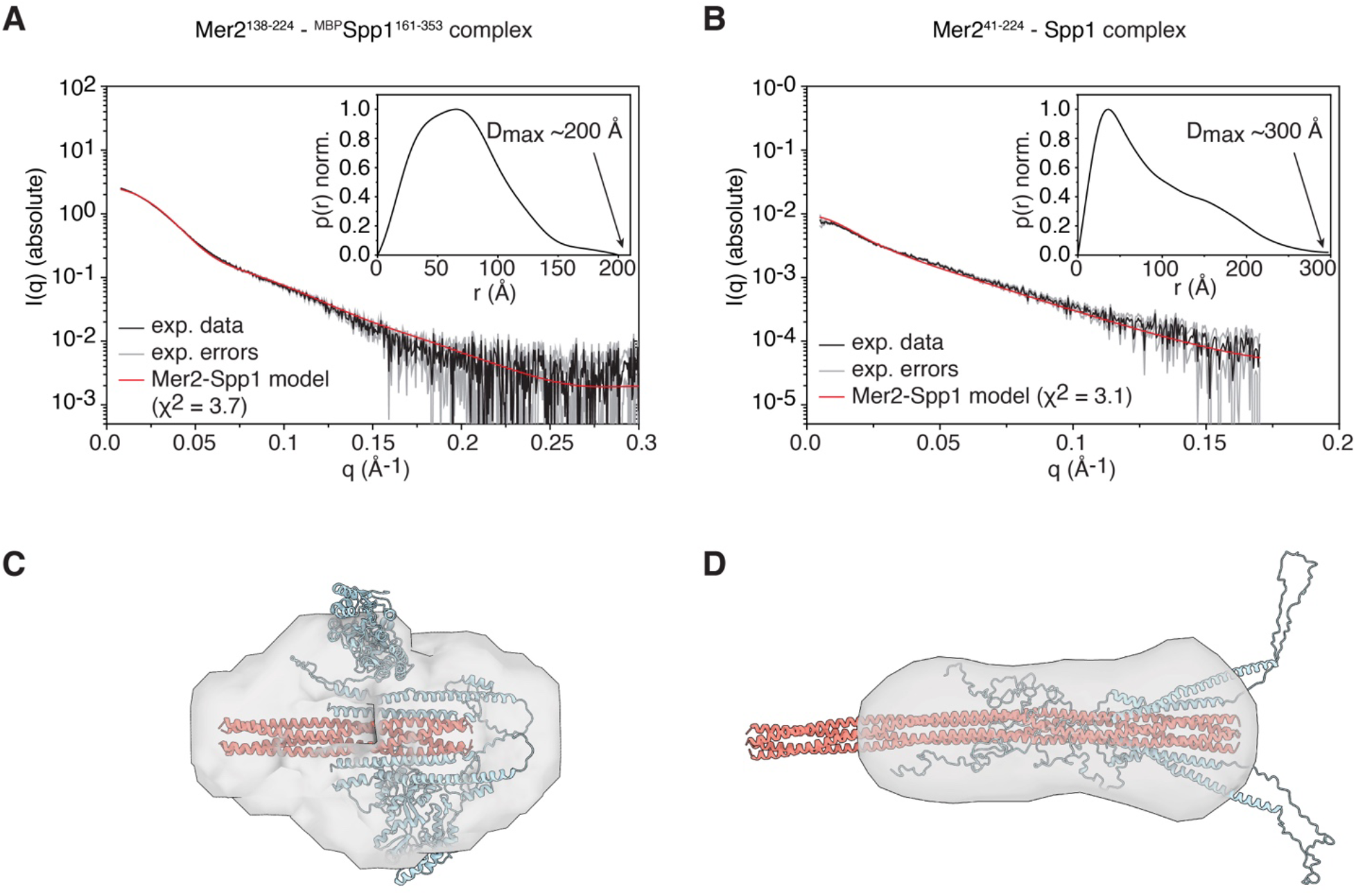
SEC-SAXS analyses of Mer2-Spp1 complexes. **(A, B)** SEC-SAXS analysis of Mer2^138-224^-^MBP^Spp1^161-353^ and Mer2^41-224^-Spp1 complexes. The main graph shows the experimental data (black), error margins (gray), and the fit of the AlphaFold model to the data (red). The inset shows a normalized probability distance distribution function obtained from the experimental SAXS data, with the approximate D_max_ value indicated. **(C, D)** Overlay of Mer2-Spp1 models with the *ab initio* reconstructed shape.

**Supplementary Figure 8:**
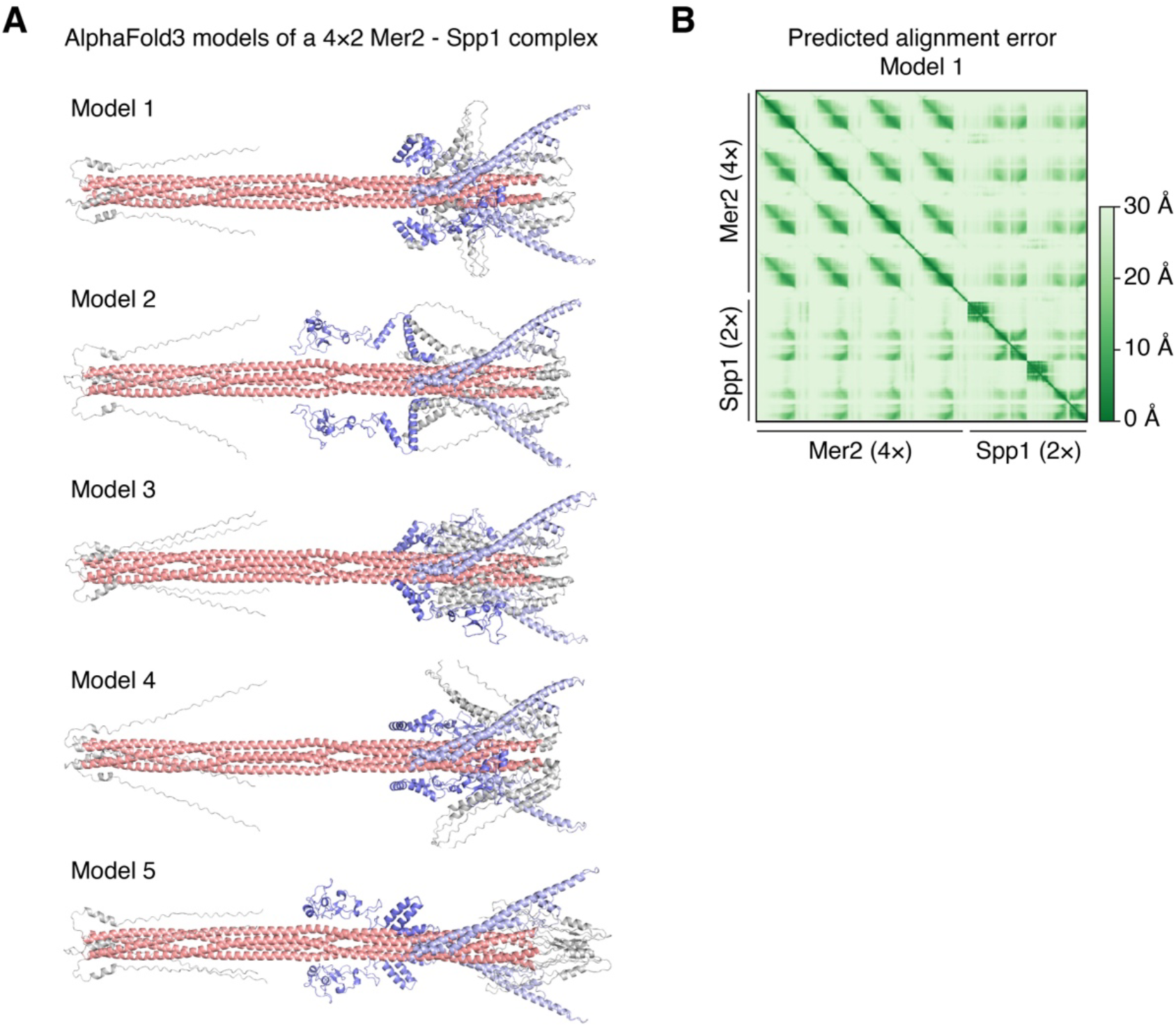
Comparisons and quality assessment of full-length Mer2-Spp1 models. **(A)** AlphaFold3 models of full-length Mer2-Spp1 complexes. The coiled coil domain of Mer2 is colored in salmon, the IDRs are grey, the N-terminal part (PHD and Set1-ID) of Spp1 is dark blue, the C-terminal part is light blue. The position of the Spp1 PHD and Set1-ID varies between models. The C-terminal Mer2 IDR is loosely placed around the Spp1-binding site. **(B)** Predicted alignment error plot for the best-scoring AlphaFold model (model 1).

**Supplementary Figure 9:**
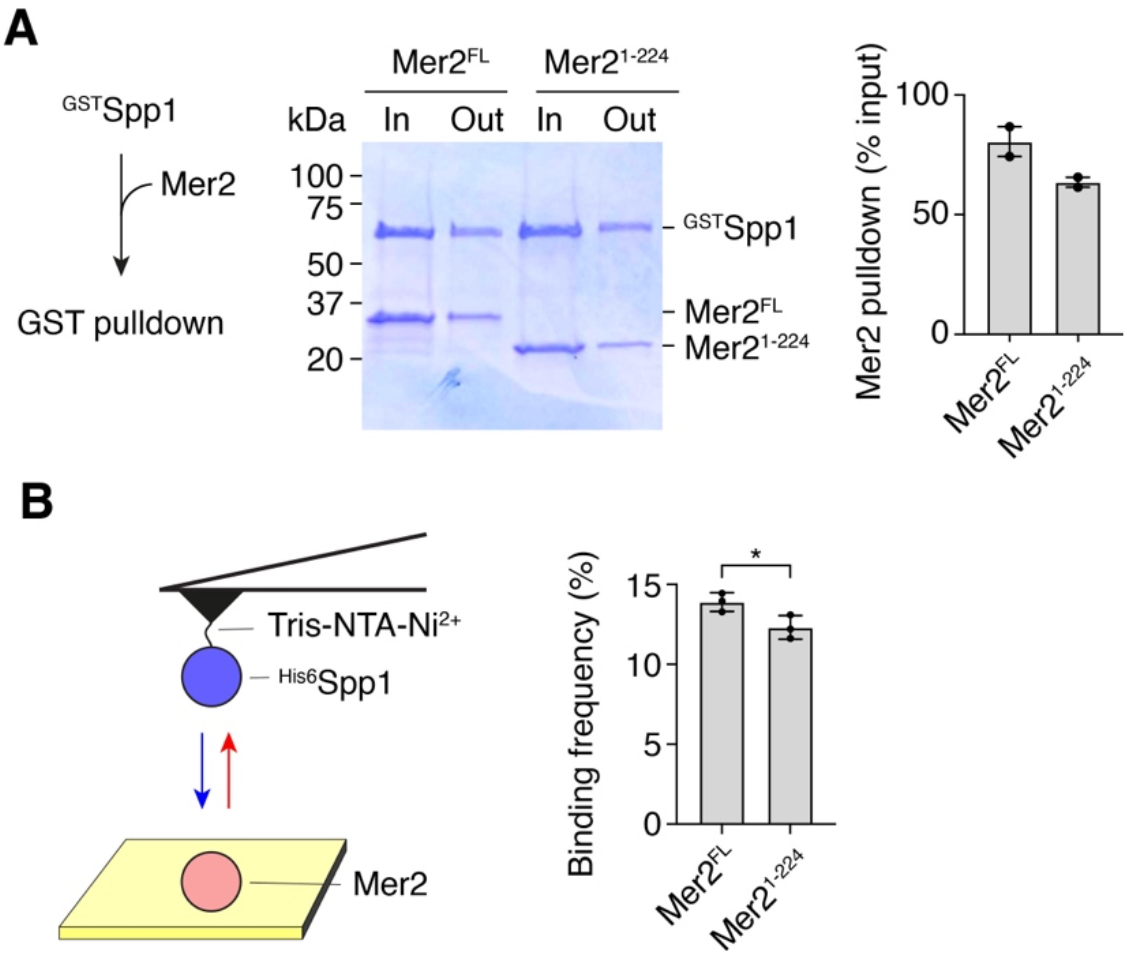
The Mer2 C-terminal IDR may weakly contribute to the interaction with Spp1. **(A)** Pulldown analysis of full-length Spp1 (2.8 µg) with full length Mer2 (1.4 µg) or a truncation of the C-terminal IDR (Mer2^1-224^) (1 µg). Error bars show the range from two independent experiments. **(B)** AFM force spectroscopy analysis of ^His^Spp1 interaction with full-length Mer2 or Mer2^1-224^. Quantification shows the binding frequency (mean + SD) from n = 3 independent experiments. *p* = 0.046 (unpaired t test).

**Supplementary Figure 10:**
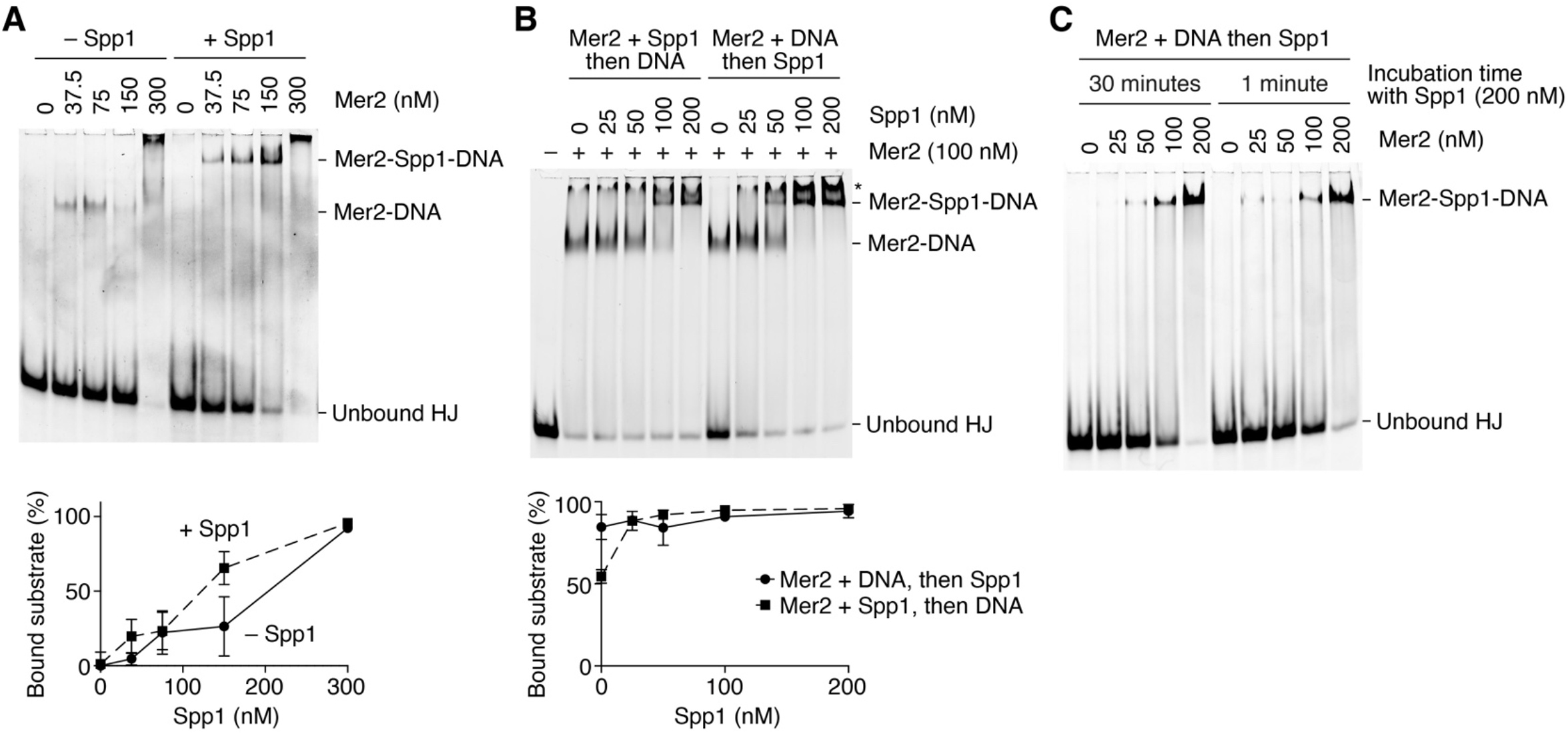
Effect of Spp1 on the DNA-binding activity of Mer2. **(A)** The effect of Spp1 (100 nM) on the binding of Mer2 to a branched DNA substrate. Quantifications show the mean ± range from two independent experiments. The fuzzy band labeled * corresponds to a higher-order Mer2-(Spp1)-DNA assembly, which we interpret as reflecting Mer2’s condensation property. The detection of this band was less reproducible, compared to the other complexes. **(B)** Gel shift analysis comparing reactions where Spp1 is added 15 minutes before (left) or after (right) Mer2 is incubated with DNA. **(C)** Gel shift analysis comparing reactions where Spp1 is added to Mer2-DNA binding reactions 30 minutes before (left) or immediately before (right) loading the gel.

**Supplementary Figure 11:**
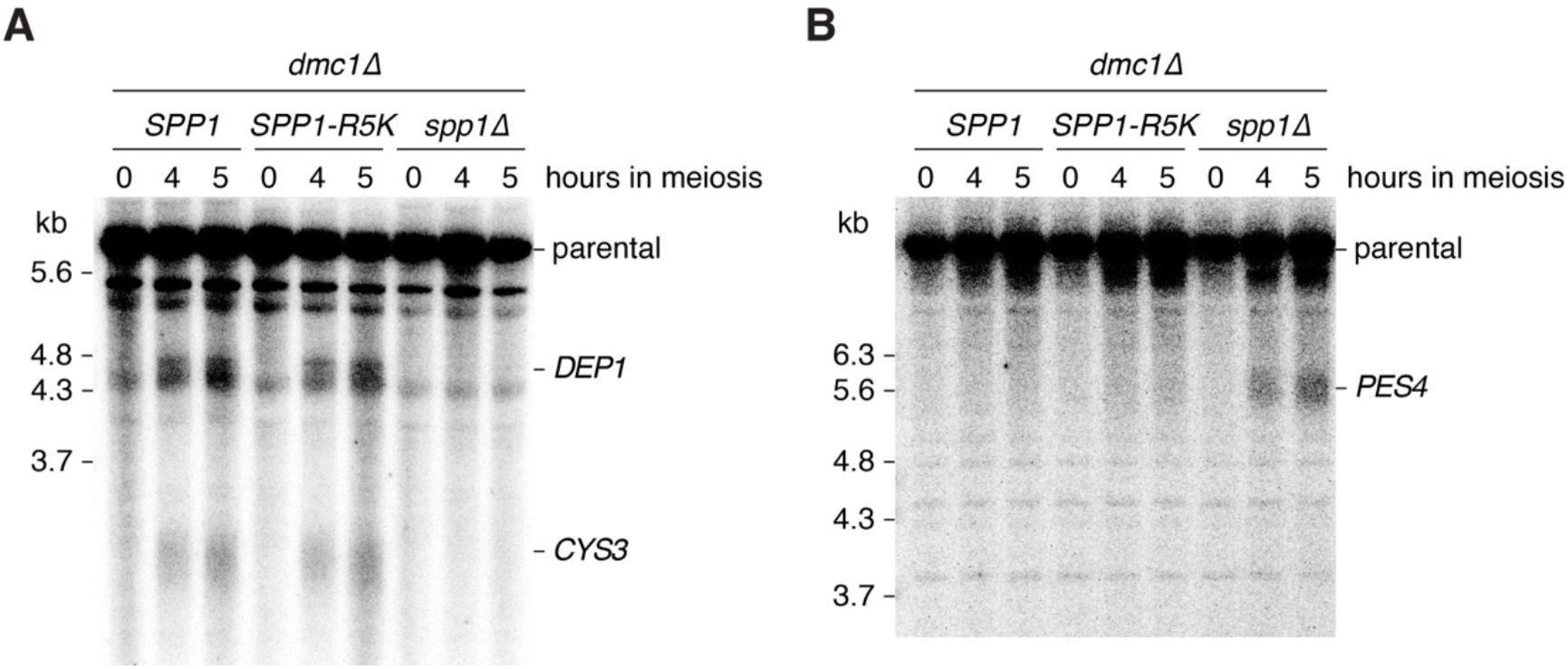
Effect of the *SPP1-R5K* mutation on meiotic DSB formation. **(A)** Southern blot analysis of meiotic DSB formation at the (A) *CYS3* and *DEP1* hotspots and (B) *PES4* hotspot in *SPP1* wild-type and mutant strains.

**Supplementary Figure 12:**
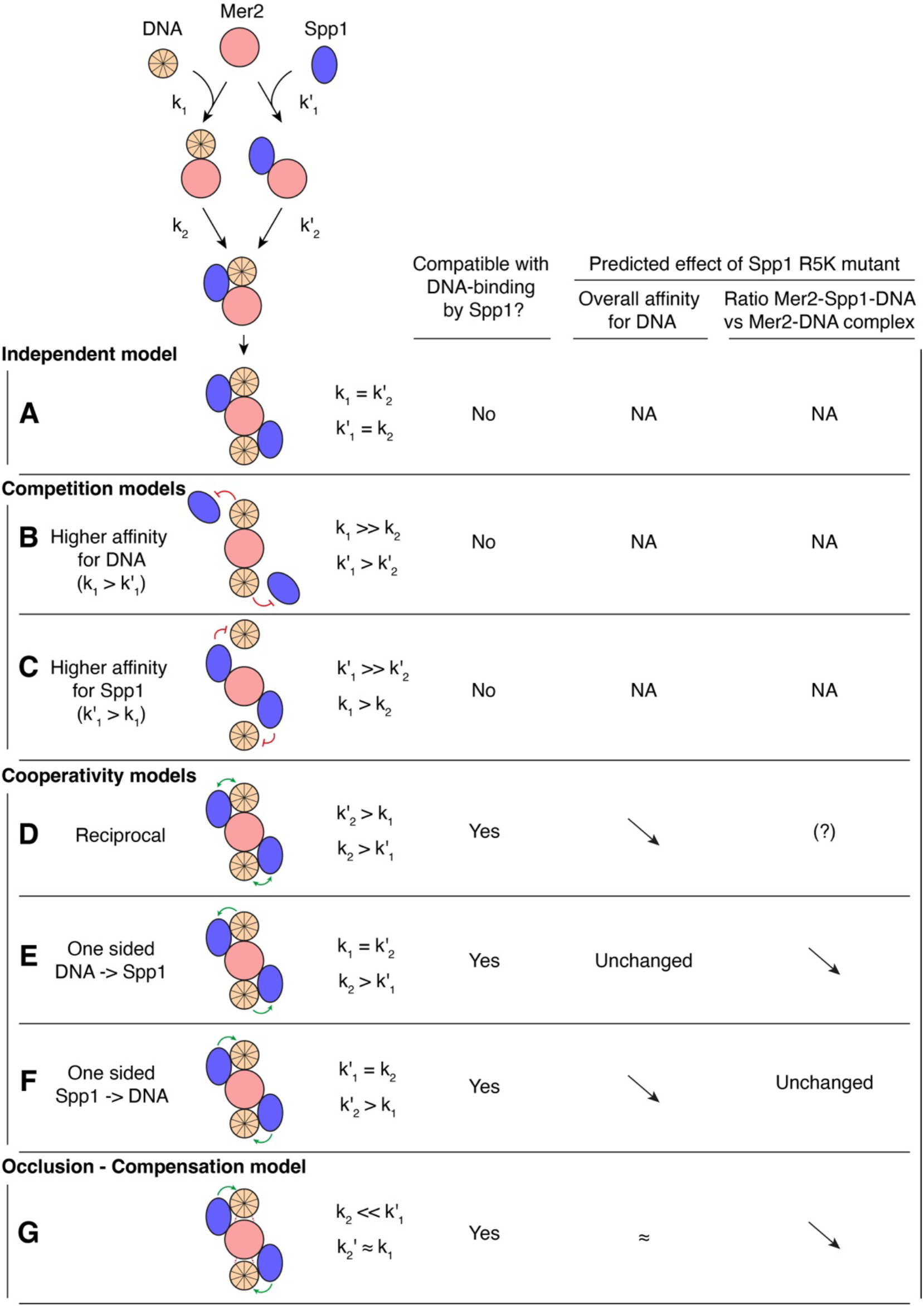
Assembly models of Mer2-Spp1-DNA complexes. Hypothetical scenarios of DNA (wheat) and Spp1 (blue) binding to the Mer2 coiled coil (pink). The cartoons illustrate a cross-section perpendicular to the coiled coil axis. The recruitment of Spp1 and DNA by Mer2 can either be independent **(A)**, competitive **(B, C)**, or cooperative **(D, E, F)**. Different scenarios can be envisioned depending on how the first binding event (k_1_) affects the second one (k_2_). The table indicates the models that are compatible with a DNA-binding activity of Spp1, and the predictions of mutating that DNA-binding activity on the affinity of the Mer2-Spp1 complex for DNA and the equilibrium between Mer2-Spp1-DNA and Mer2-DNA complexes. Based on those predictions, our results are mostly compatible with the occlusion-compensation model **(G)**, where Spp1 reduces DNA-binding by Mer2 but compensates this decreased activity by providing an additional DNA-binding site.

**Supplementary Table 1:** AFM analysis of Mer2-Spp1 association and dissociation rates.

| Model | $k_{on}$ |
| --- | --- |
| Equation | $BF = A * \left[ 1 - \exp\left(-\frac{t-t_0}{\tau}\right) \right]$ |
| $\tau$ | $75 \pm 12$ ms |
| $t_0$ | $-3.1 \pm 6.4$ ms |
| A | $13.6 \pm 0.71$ % |
| $\chi^2$ | 1.48 |
| $R^2$ | 0.96 |

| Model | Bell-Evans fit |
| --- | --- |
| Equation | $F(r) = \left( \frac{k_B T}{x_\beta} \right) \ln \frac{r x_\beta}{k_B T k_{off}}$ |
| $x_\beta$ | $0.76 \pm 0.06$ nm |
| $k_{off}$ | $0.19 \pm 0.15$ s<sup>-1</sup> |
| $\chi^2$ | 0.14 |
| $R^2$ | 0.89 |

**Supplementary Table 2:**
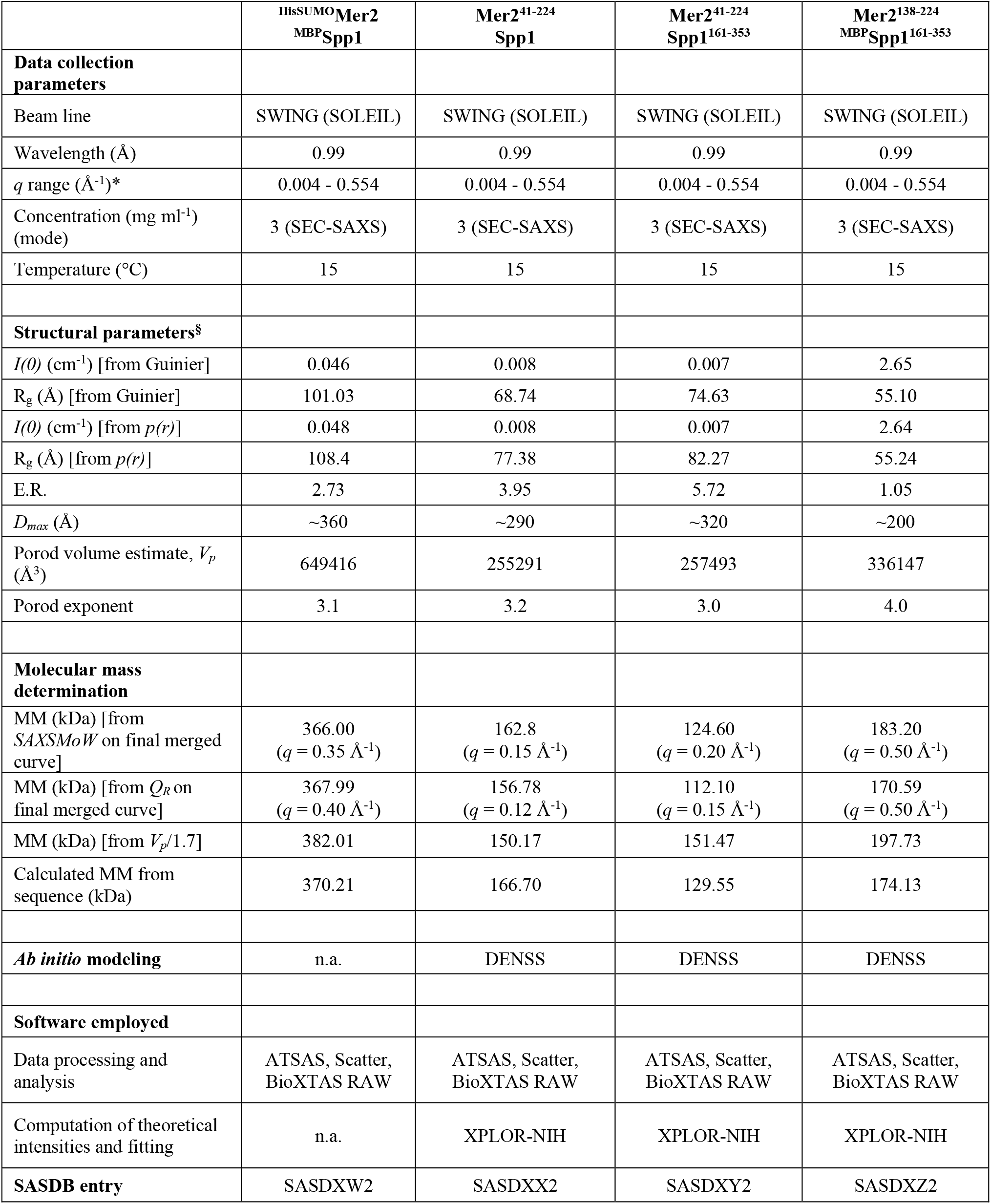
SAXS data collection and scattering-derived parameters. Abbreviations: *I(0)*, extrapolated scattering intensity at zero angle; *R*_*g*_, radius of gyration calculated using either Guinier approximation (from Guinier) or the indirect Fourier transform package GNOM [from *p(r)*]; *MM*, molecular mass; *D*_*max*_, maximal particle dimension; *V*_*p*_, Porod volume; E.R., elongation ratio. *Momentum transfer |*q*| = 4πsin(θ)/λ.

**Supplementary Table 3:** Plasmids used in this study.

| Plasmid | Description | Reference |
| --- | --- | --- |
| pPL001 | HisSUMO-Spp1 in pSMT3 | This study |
| pPL011 | MBP-Tev-Mer2 in pMAL-c2x | This study |
| pPL012 | MBP-Tev-Spp1 in pMAL-c2x | This study |
| pPL014 | HisSUMO-Mer2 WT, MBP-Tev-Spp1 WT in pEtDuet-1 | This study |
| pPL015 | GST in pEtDuet-1 | This study |
| pPL016 | GST-Tev-Spp1 WT in pEtDuet-1 | This study |
| pPL032 | GST-Tev-mScarlet-Spp1 WT in pCCB205 | This study |
| pPL036 | MBP-Tev-Mer2 V195D in pMAL-c2x | This study |
| pPL037 | HisSUMO-eGFP-Mer2 V195D in pSMT3 | This study |
| pPL041 | HisSUMO-Mer2 (138-224) in pSMT3 | This study |
| pPL042 | MBP-Spp1 (161-353) in pMAL-c2x | This study |
| pPL045 | HisSUMO-Mer2 (138-224) in pEtDuet-1 | This study |
| pPL053 | MBP-Spp1 (161-353) in pEtDuet-1 | This study |
| pPL054 | HisSUMO-Mer2 (138-224) WT, MBP-Tev-Spp1 (161-353) in pEtDuet-1 | This study |
| pPL064 | HisSUMO-Mer2 (138-224) WT, MBP-Tev-Spp1 (161-353) Q317A in pEtDuet-1 | This study |
| pPL065 | HisSUMO-Mer2 (138-224) WT, MBP-Tev-Spp1 (161-353) Q318A in pEtDuet-1 | This study |
| pPL066 | HisSUMO-Mer2 (138-224) WT, MBP-Tev-Spp1 (161-353) S324A in pEtDuet-1 | This study |
| pPL067 | HisSUMO-Mer2 (138-224) WT, MBP-Tev-Spp1 (161-353) R328A in pEtDuet-1 | This study |
| pPL068 | HisSUMO-Mer2 (138-224) WT, MBP-Tev-Spp1 (161-353) L332A in pEtDuet-1 | This study |
| pPL069 | HisSUMO-Mer2 (138-224) WT, MBP-Tev-Spp1 (161-353) R336A in pEtDuet-1 | This study |
| pPL070 | HisSUMO-Mer2 (138-224) WT, MBP-Tev-Spp1 (161-353) Q339A in pEtDuet-1 | This study |
| pPL071 | HisSUMO-Mer2 (138-224) WT, MBP-Tev-Spp1 (161-353) I342A in pEtDuet-1 | This study |
| pPL072 | HisSUMO-Mer2 (138-224) N184A, MBP-Tev-Spp1 (161-353) WT in pEtDuet-1 | This study |
| pPL073 | HisSUMO-Mer2 (138-224) Q188A, MBP-Tev-Spp1 (161-353) WT in pEtDuet-1 | This study |
| pPL074 | HisSUMO-Mer2 (138-224) E190A, MBP-Tev-Spp1 (161-353) WT in pEtDuet-1 | This study |
| pPL075 | HisSUMO-Mer2 (138-224) V195D, MBP-Tev-Spp1 (161-353) WT in pEtDuet-1 | This study |
| pPL076 | HisSUMO-Mer2 (138-224) D202A, MBP-Tev-Spp1 (161-353) WT in pEtDuet-1 | This study |
| pPL077 | HisSUMO-Mer2 (138-224) T208A, MBP-Tev-Spp1 (161-353) WT in pEtDuet-1 | This study |
| pPL078 | HisSUMO-Mer2 (138-224) R209A, MBP-Tev-Spp1 (161-353) WT in pEtDuet-1 | This study |
| pPL079 | HisSUMO-Mer2 (138-224) E214A, MBP-Tev-Spp1 (161-353) WT in pEtDuet-1 | This study |
| pPL089 | HisSUMO-Mer2 (138-224) V195A, MBP-Tev-Spp1 (161-353) WT in pEtDuet-1 | This study |
| pPL090 | HisSUMO-Mer2 (138-224) WT, MBP-Tev-Spp1 (307-353) WT in pEtDuet-1 | This study |
| pPL094 | GST-Tev-Spp1 (161-353) WT in pEtDuet-1 | This study |
| pPL095 | GST-Tev-Spp1 (161-353) R5K in pEtDuet-1 | This study |
| pPL099 | GST-Tev-Spp1 R5K in pEtDuet-1 | This study |
| pCCB205 | pMAL-c2x | New England Biolabs |
| pCCB407 | pFastBac1-GST | This study |
| pCCB750 | HisSUMO-Mer2 WT in pSMT3 | Claeys Bouuaert et al., 2021 |
| pCCB777 | HisSUMO-eGFP-Mer2 WT in pSMT3 | Claeys Bouuaert et al., 2021 |
| pCCB785 | HisSUMO-mScarlet-Mer2 WT in pSMT3 | This study |
| pDD057 | HisSUMO-Mer2 (41-314) in pSMT3 | Daccache et al., 2023 |
| pDD060 | HisSUMO-Mer2 (1-224) in pSMT3 | Daccache et al., 2023 |
| pDD065 | HisSUMO-Mer2 (41-224) in pSMT3 | Daccache et al., 2023 |
| pKM001 | His-Mer2 WT in pET28b | This study |
| pKM004 | GST-His-Spp1 WT in pEtDuet-1 | This study |

**Supplementary Table 4:**
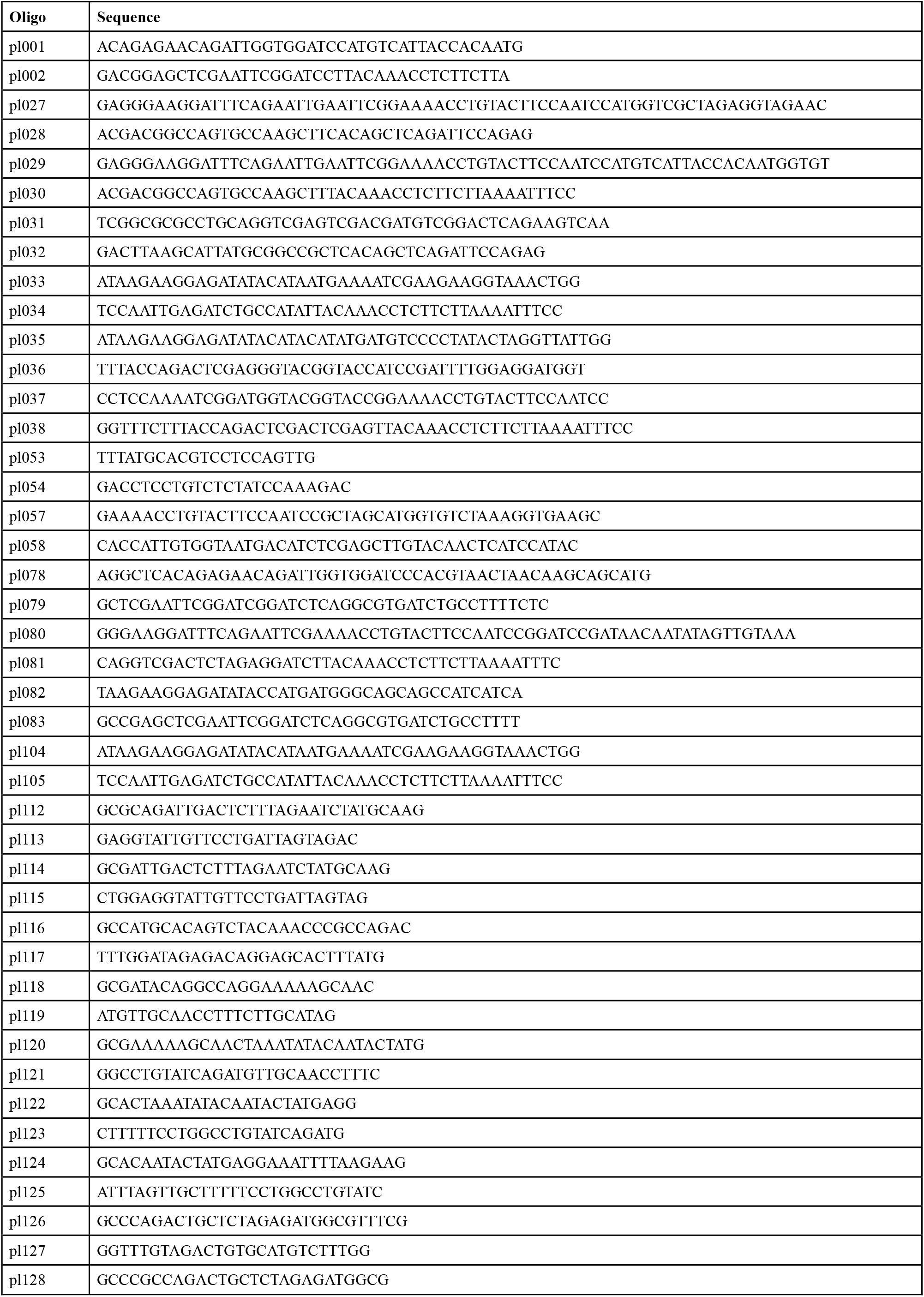
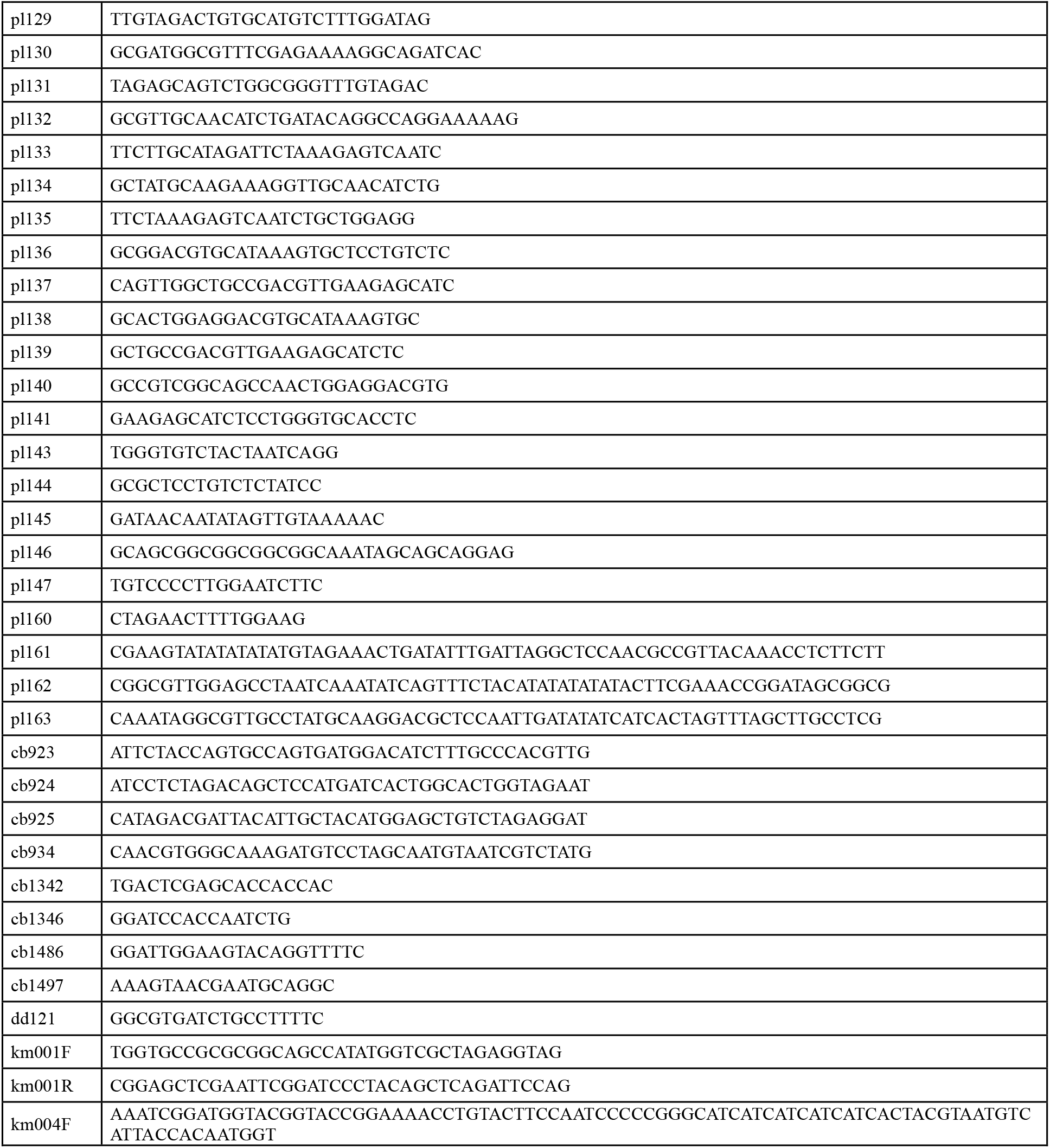
Oligonucleotides used in this study.

**Supplementary Table 5:**
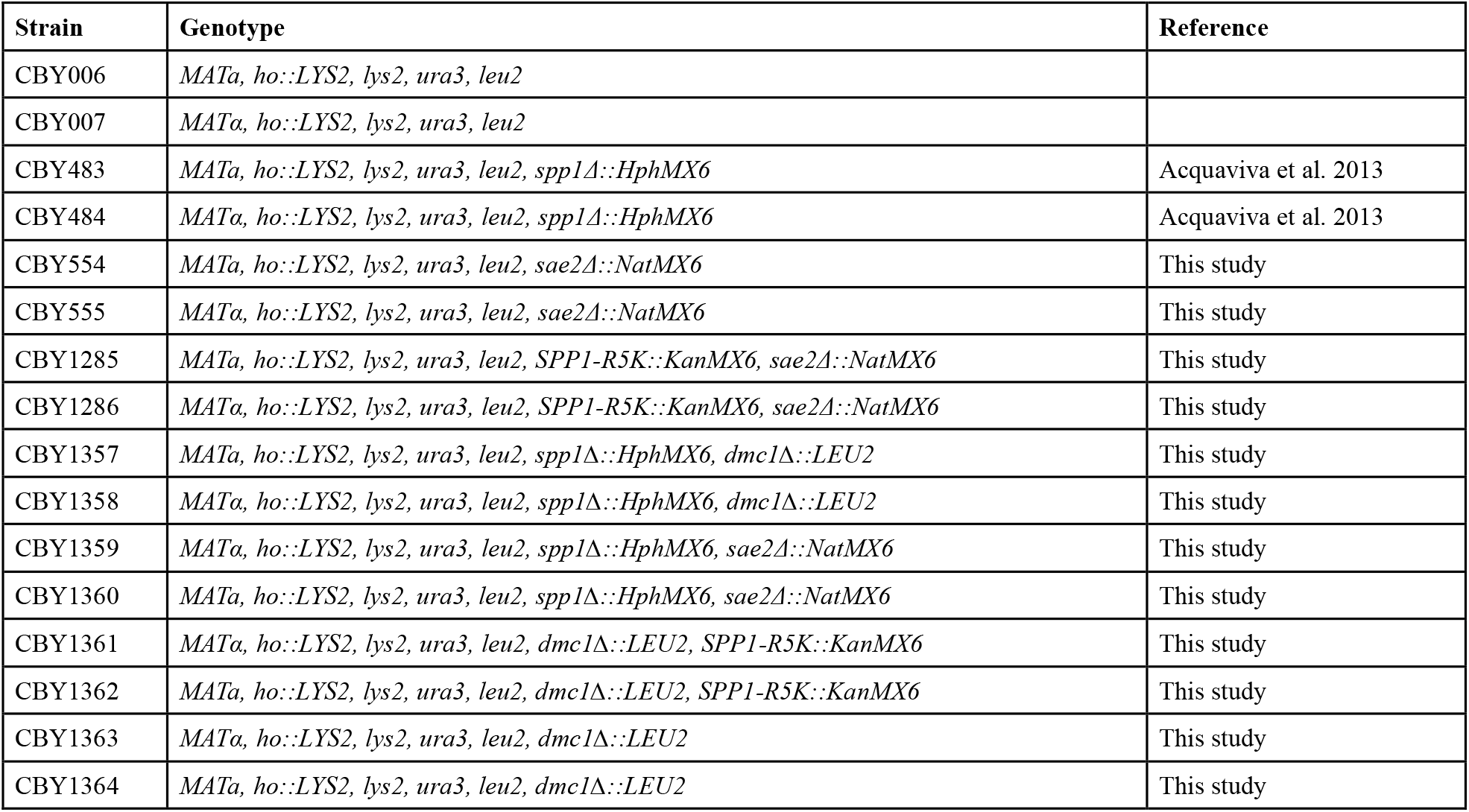
Yeast strains used in this study. All strains are from the SK1 background.

**Supplementary Table 6:**
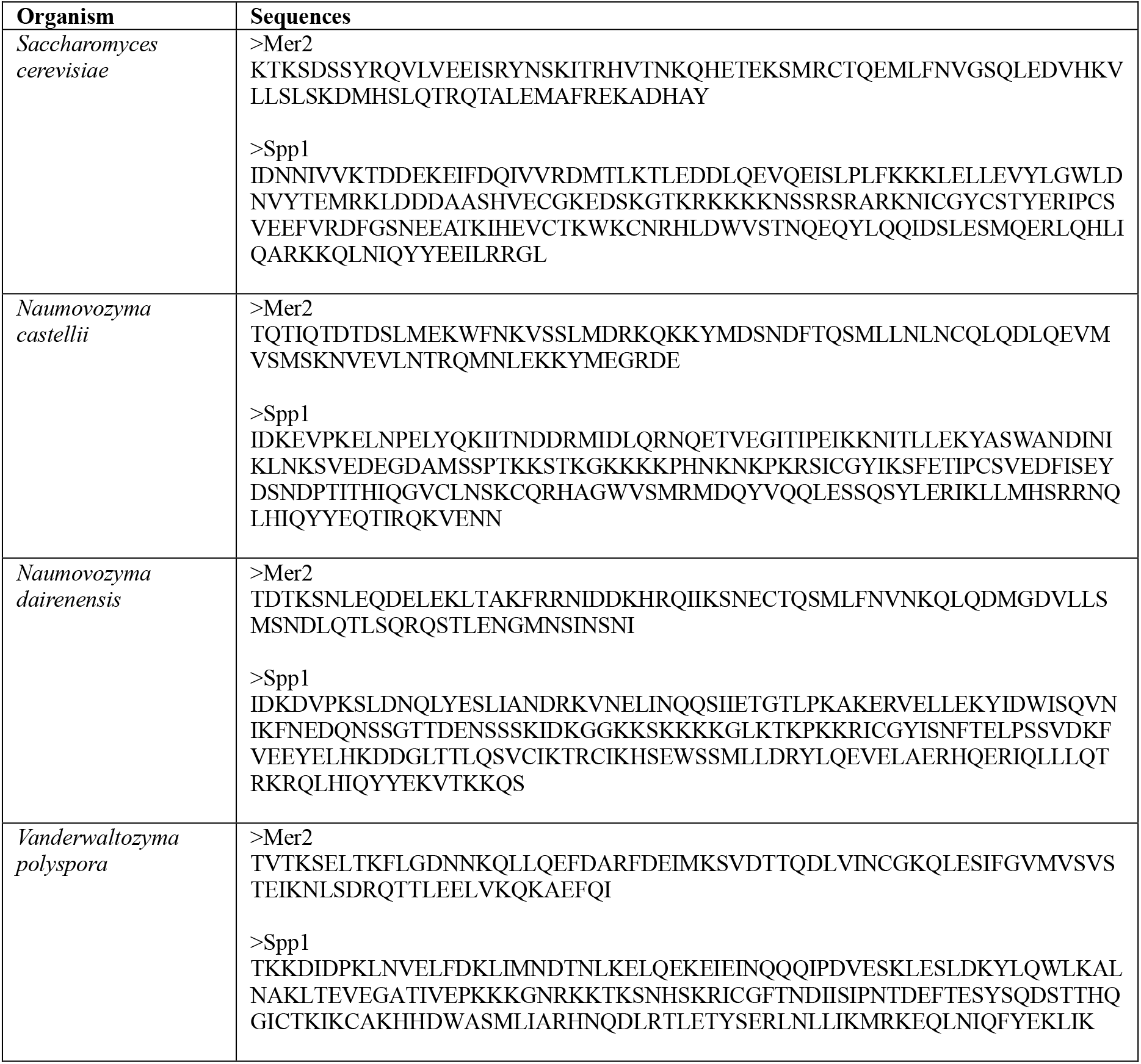
Input sequences for AlphaFold3 modeling. Sequences used to generate models presented in **Figure 2A, Supplementary Figure 3A**, and **Supplementary Figure 4**.

